# Mechanical Responses of the Giant Nesprin-2

**DOI:** 10.1101/2025.05.30.657030

**Authors:** Fei Shang, Yuhang Zhang, Jiaqing Ye, Zhuwei Zhang, Xingyu Qi, Hu Chen, Miao Yu, Shimin Le

## Abstract

The nesprin protein family serves as a critical physical bridge between the cytoskeleton–a fundamental structural scaffold and mechanotransduction hub of the cell, and the nucleus–an intriguing and emerging mechano-responsive element. Due to the external mechanical cues and the nucleo-cytoskeletal dynamics, the nesprins are physiologically under forces. However, the dynamics of nesprins within physiological forces and loading rates remain largely unexplored. In this study, we employ magnetic tweezers based single-molecule manipulation alongside molecular dynamic simulations and AlphaFold structural predictions to comprehensively investigate the dynamics of force-bearing spectrin repeat (SR) domains of the giant nesprin-2 protein. Through direct quantification, we unveil that the numerous SRs undergo mechanical unfolding and refolding dynamics with distinct transition rates within several pN scale. Furthermore, we show that the giant nesprin-2 could act as an effective molecular absorber adeptly maintaining forces on the nucleoskeleton and cytoskeleton linkage within a few pN across displacement spans exceeding ***µ***m. Notably, our findings imply that subtle pN-level mechanical forces intricately modulate nesprin–protein interactions via the dynamics of domain folding and unfolding. Collectively, our study offers a comprehensive understanding of the mechanical characteristics of giant nesprin-2, shedding light on its pivotal role in nucleoskeleton–cytoskeleton mechanotransduction.

## Introduction

Cell cytoskeleton, the major structural framework and mechanoresponsive element, is a dynamic and adaptable network comprising interconnecting protein filaments, namely, actin filaments, intermediate filaments, and microtubules, along with associated motor proteins, adaptor proteins, and cross-linking proteins [1–4]. While the cell cytoskeleton mediated mechanobiology has garnered significant attention over recent decades, particularly in proximity to the cell membrane and the extracellular microenvironment [1–4], a substantial gap persists in our understanding of how the cell’s cytoskeleton engages in mechanosensing, mechanotransmission, and mechanotransduction with the nucleus-a multiple-role organelle serving as a genetic repository and transcription site[5–8]. As the nucleus emerges as an intriguing mechanoresponsive element, an in-depth exploration of its mechanical interplay with the cytoskeleton becomes a compelling frontier in the research of cell mechanobiology[5–8].

The pivotal physical supramolecular link between the cytoskeleton and the nucleus, essential for nucleocytoskeletal coupling, is established through the LINC (linker of the nucleoskeleton and the cytoskeleton) complex [5–8]. LINC complex comprises the cytoplasmic nesprin protein family and the inner nuclear membrane (INM) located SUN (Sad1p and UNC-84 homology) proteins[9, 10]. The critical importance of nesprins is underscored by the fact that mutations or protein loss results in various detrimental cellular effects, including impaired nuclear envelope integrity [11], compromised nucleus anchoring [12, 13], disrupted signaling to the nucleus [14], altered chromosome positioning [15], hindered DNA repair [16], perturbed genome transcription [17], hindered replication and transcription [18–21], as well as impaired response to extracellular mechanical cues [22]. Moreover, mutations or dysregulation of nesprins have been associated with pathologic conditions such as muscular dystrophies, cardiomyopathies, and neurodegenerative disorders[23–27].

The nesprins are anchored to the nuclear envelope via the interaction of its C-terminal short KASH (Klarsicht, ANC-1, and Syne homology) domain with the the C-terminal region of SUN proteins within the perinuclear space [9, 10], thereby also indirectly associated with the nuclear lamina, the nuclear pore complex (NPCs), and chromatin[28, 29]. The N-terminal region of nesprins, a pair of calponin homology domains, represents a classic actin-binding domain (ABD), reminiscent of *α*-actinin, dystrophin, and plectin etc[3, 30, 31], enabling nesprins directly linking to actin cytoskeleton. Moreover, nesprin-1, nesprin-2 and nesprin-4 can associate with microtubules through interactions with kinesin and dynein [32–34], while nesprin-3 physically connects to intermediate filament vimentin via the plectin protein [35]. Thereby, the nesprins physically connect all three major cytoskeletal filaments to the nucleus. Bridging the KASH and ABD domains are a notable number spectrin repeats(SRs) in the giant nesprin-1 (74 SRs) and nesprin-2 (56 SRs), as well as nesprin-3 (9 SRs), respectively[32, 36, 37]. Therefore, the numerous SRs on nesprins that connect the cytoskeleton and nucleus are physiologically under forces.

Notably, recent fluorescence resonance energy transfer(FRET) tension sensor or helix-hairpin-based tension senor studies on a truncated nesprin-2 (mini-nesprin-2G) has demonstrated that nesprin is subjected to cytoskeleton-dependent tension in pN scale, and is sensitive to extracellular mechanical cues [37–40]. Tumor cells enable extensive deformation of the cell and its nucleus in order to penetrate tissues through tight interstitial spaces during cancer metastasis[41]. During such confined cell migration, nesprin-2 accumulates at the front of the nucleus, which is regulated by the mechanical forces resulted from actin cytoskeleton [42]. However, a key question that how nesprins, particularly the giant nesprins, response to the physiological level forces is still unexplored.

In this study, we employ magnetic-tweezers-based single-molecule manipulation alongside molecular dynamic simulations and AlphaFold2 structural prediction and analysis to comprehensively investigate the responses of the force-bearing structural domains of the giant nesprin-2 protein within physiologically relevant force range and force loading rates. Through direct quantification, we unveil that all 56 SRs undergo mechanical unfolding at forces below 25 pN. Furthermore, we show that the giant nesprin-2 could at as a molecular absorber adeptly maintaining forces on the nucleoskeleton and cytoskeleton linkage within the range of 5 pN across displacement spans exceeding *µ*m. Notably, our findings emphasize that subtle pN-level mechanical forces intricately modulate nesprin-protein interactions via the dynamics of domain folding and unfolding. Collectively, our study offers a comprehensive understanding of nesprin-2’s mechanical characteristics, shedding light on its pivotal role in nucleo-cytoskeleton mechanotransduction.

## Results

### Responses of the giant nesprin-2 over physiological force-loading

To systematically characterize the force responses of the giant nesprin-2 protein, we divided all 56 SRs of giant nesprin-2 into 26 single-molecule constructs based on the UniProt database (accession Q6ZWQ0-1) and the AlphaFold2 predicted structure models[43] (Fig. 1a-b, see Methods: Plasmids and proteins). In each construct, one to three neighboring SR domains are positioned between an N-terminal biotinylated AviTag and a C-terminal SpyTag. These two tags enable specifically tethering of the target protein domains between a streptavidin-coated superparamagnetic bead and a SpyCatcher-coated coverslip surface in a flow channel [44–46]. Subsequently, we applied well-controlled external force to the tethered protein domains using a magnetic tweezers setup [47, 48] and observed the real-time structural dynamics of the domains under force, reflected by corresponding changes in the tethered bead height along the force direction (Fig. 1c-d, see Methods: Magnetic tweezers manipulation).

**Fig. 1.**
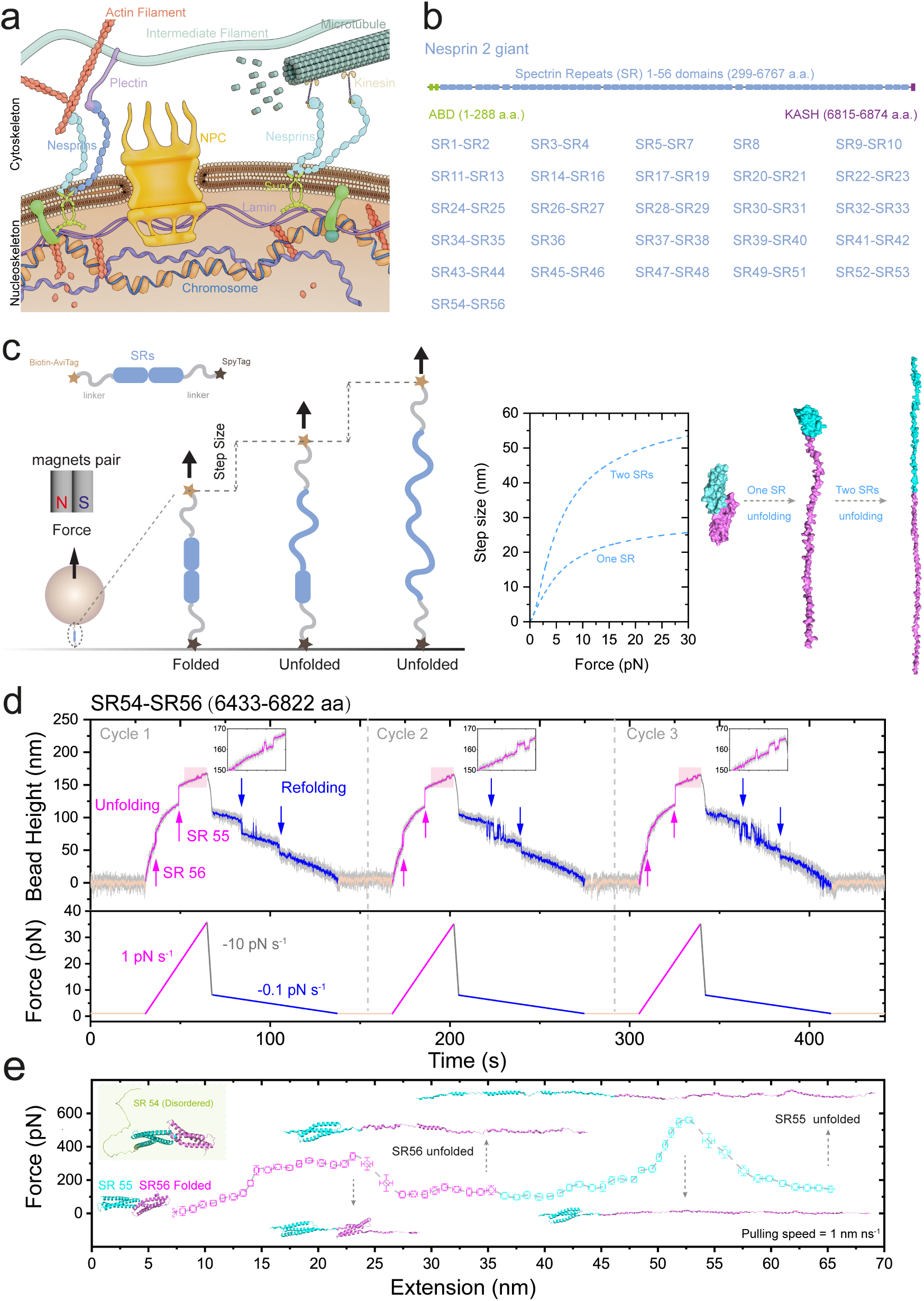
Single-molecule manipulation on giant nesprin-2. (**a**) Schematic of the nesprin mediated mechanopathways at the outer and inter nucleomembrane. (**b**) Illustrations of single-molecule constructs of giant nesprin-2. Bottom panel: segmentalization of the giant nesprin-2. (**c**) Left panel: illustrations of single-molecule manipulation. Right panel: theoretical force-dependent unfolding/refolding step sizes of one or two SR domains with structural illustrations based on AlphaFold2 predicted SR55-SR56. (**d**) Representative cycles of time–bead height curves of SRs (e.g., SR54-SR56) during linear force- increase (magenta) and force-decrease (blue) scans with a loading rate of 1 pN s^-1^ and -0.1 pN s^-1^, respectively. The raw data (gray) is 10-point Fast Fourier Transform (FFT) smoothed (colored lines). Force-dependent unfolding and refolding events are indicated by magenta upward arrows and blue downward arrows, respectively. Insets: the zoom-in of SpyTag unzipping/zipping dynamics at ∼30 pN. (**e**) Representative extension–force curve of SR55-SR56 unfolding dynamics obtained in steered molecular dynamics simulations with constant pulling at 1 nm ns^-1^). Snaps of the structural dynamics of the SR55-SR56 are plotted as insets. The Left-top inset shows the AlphaFold2 predicted structure of SR54-SR56.

Figure 1d shows an example of force-loading scan cycles for one of the constructs (SR54-SR56 at the C-terminus), where the force was linearly increased at a physiologically relevant loading rate of 1 pN s^-1^ [49, 50] to probe the unfolding of the SR domains, indicated by the force-dependent unfolding steps (magenta arrows in the panel). Once the domain unfolded, the force was rapidly decreased (at a loading rate of -10 pN s^-1^) to ∼8 pN, and then switched to a slow loading rate of -0.1 pN s^-1^ to ∼1 pN to observe the refolding of the SR domains under low forces (blue arrows in Fig. 1d). In addition to the conformational dynamics of the target SR domains, rapid unfolding and refolding fluctuations with a step size of ∼5 nm observed at ∼30 pN were due to the unzipping and zipping of the SpyTag from SpyCatcher (zoomed-in insets in Fig. 1d and Supplementary Fig. 1). This specific signal serves as a fingerprint of the specific tethering of the target molecules, as well as an additional force-calibration reference.

Interestingly, for the SR54-SR56 construct, only two unfolding events corresponding to the unfolding of two SR domains were observed. AlphaFold2 structural prediction for SR54-56 (as well as SR52-SR56) suggests that SR54 is in an unstructured state in the absence of force (Fig. 1d, Supplementary Fig. 6 and Supplementary Fig. 7b, See Methods: AlphaFold structural prediction)[43]. Consistent with this structural prediction, the single-molecule manipulation of SR54-SR56 leads to two unfolding and refolding events, corresponding to the structural dynamics of SR55 and SR56. These combined findings provide compelling evidence that SR54 exists as an intrinsically disordered domain under physiological conditions.

Since each single-molecule construct contains one to three SR domains, we precisely assigned the mechanical responses to each domain by performing steered molecular dynamics (SMD) simulations on AlphaFold2-predicted structures for each construct, enabling differentiation of their distinct mechanical behaviors (see Methods: AlphaFold structural prediction, Steered molecular dynamics simulations)[43]. Figure 1e displays representative structural snaps of the SR54-SR56 domains during constant pulling (1 nm s^-1^) in SMD, capturing conformations before and after the unfolding transitions. These simulations clearly demonstrate that the SR55 domain is mechanically stronger than SR56; consequently, we attributed the more mechanically robust responses to SR55 and the less stable responses to SR56. Complete structural snapshots for all SR domains are presented in panels i of Fig. 2-5, with full dynamic trajectories available in Supplementary Mov. 1-26.

**Fig. 2.**
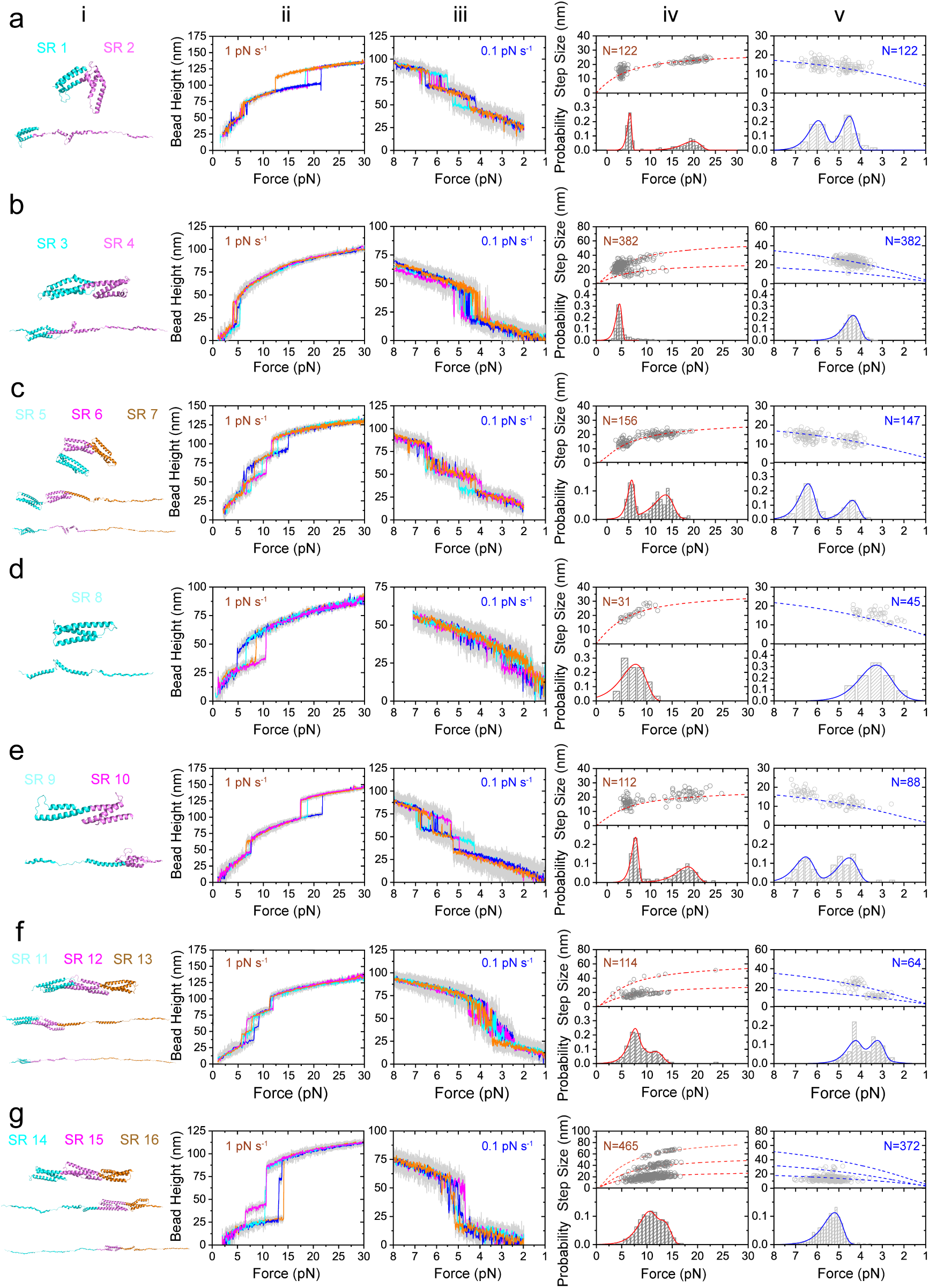
Force responses of giant nesprin-2 SR1-SR16 during linear force-loading. (**a-g**) Panel i: Representative snaps of corresponding SR domains before and after unfolding transition extracted from SMD simulations on AlphaFold2 predicted structures. Panels ii and iii: Representative force–bead height curves of the corresponding SR domains during linear force-increase (ii, 1 pN s^-1^) and force-decrease (iii, -0.1 pN s^-1^) scans. Raw data (gray) is 10-point FFT smoothed (colored lines). Panels iv and v: The resulting force–step size distributions of unfolding events (top, iv) and refolding events (top, v) obtained from the linear scans. The resulting normalized unfolding force (bottom, iv) and refolding force (bottom, v) distributions of the SRs. Number (*N*) of data points (represented by hollow circles) are indicated in the panels. The dashed colored lines are theoretical calculation of the force-dependent unfolding/refolding step sizes for corresponding SR domains. The colored solid lines are fitted force distributions based on Bell’s model (unfolding) or Arrhenius Law (refolding).

**Fig. 3.**
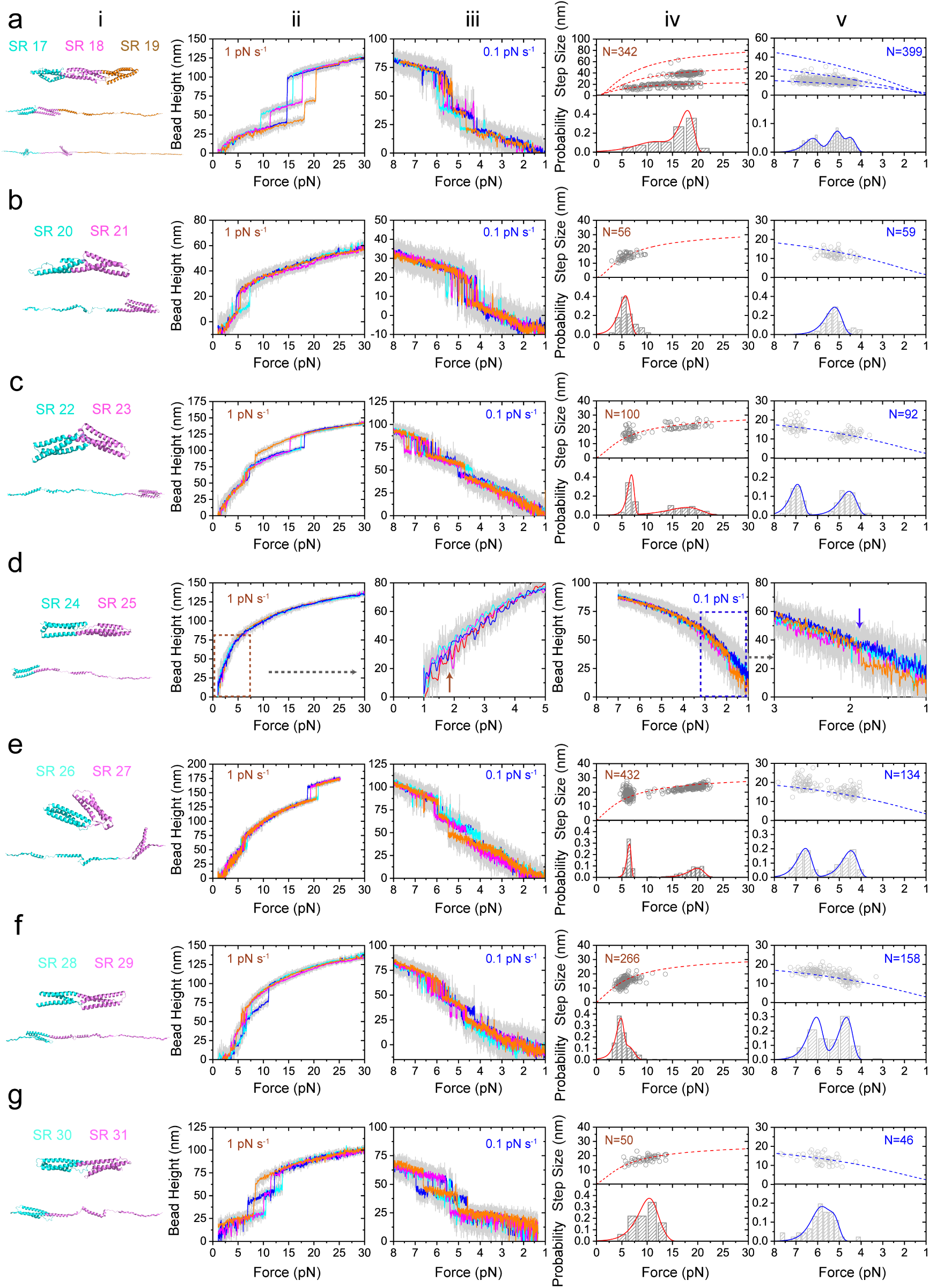
Force responses of giant nesprin-2 SR17-SR31 during linear force-loading. (**a-c,e-g**) Panel i: Representative snaps of corresponding SR domains before and after unfolding transition extracted from SMD simulations on AlphaFold2 predicted structures. Panels ii and iii: Representative force–bead height curves of the corresponding SR domains during linear force-increase (ii, 1 pN s^-1^) and force-decrease (iii, -0.1 pN s^-1^) scans. Raw data (gray) is 10-point FFT smoothed (colored lines). Panels iv and v: The resulting force–step size distributions of unfolding events (top, iv) and refolding events (top, v) obtained from the linear scans. The resulting normalized unfolding force (bottom, iv) and refolding force (bottom, v) distributions of the SRs. Number (*N*) of data points (represented by hollow circles) are indicated in the panels. The dashed colored lines are theoretical calculation of the force-dependent unfolding/refolding step sizes for corresponding SR domains. The colored solid lines are fitted force distributions based on Bell’s model (unfolding) or Arrhenius Law (refolding). For SR20-SR21, only one unfolding/refolding event (SR21) in each cycle was observed. (**d**) For SR24-SR25, clear unfolding/refolding events was not observed. Panel iii and v are zoom-in of the force–bead height curves of ii and iv at low force regime, respectively. Arrows in iii and v indicate potential transition events.

**Fig. 4.**
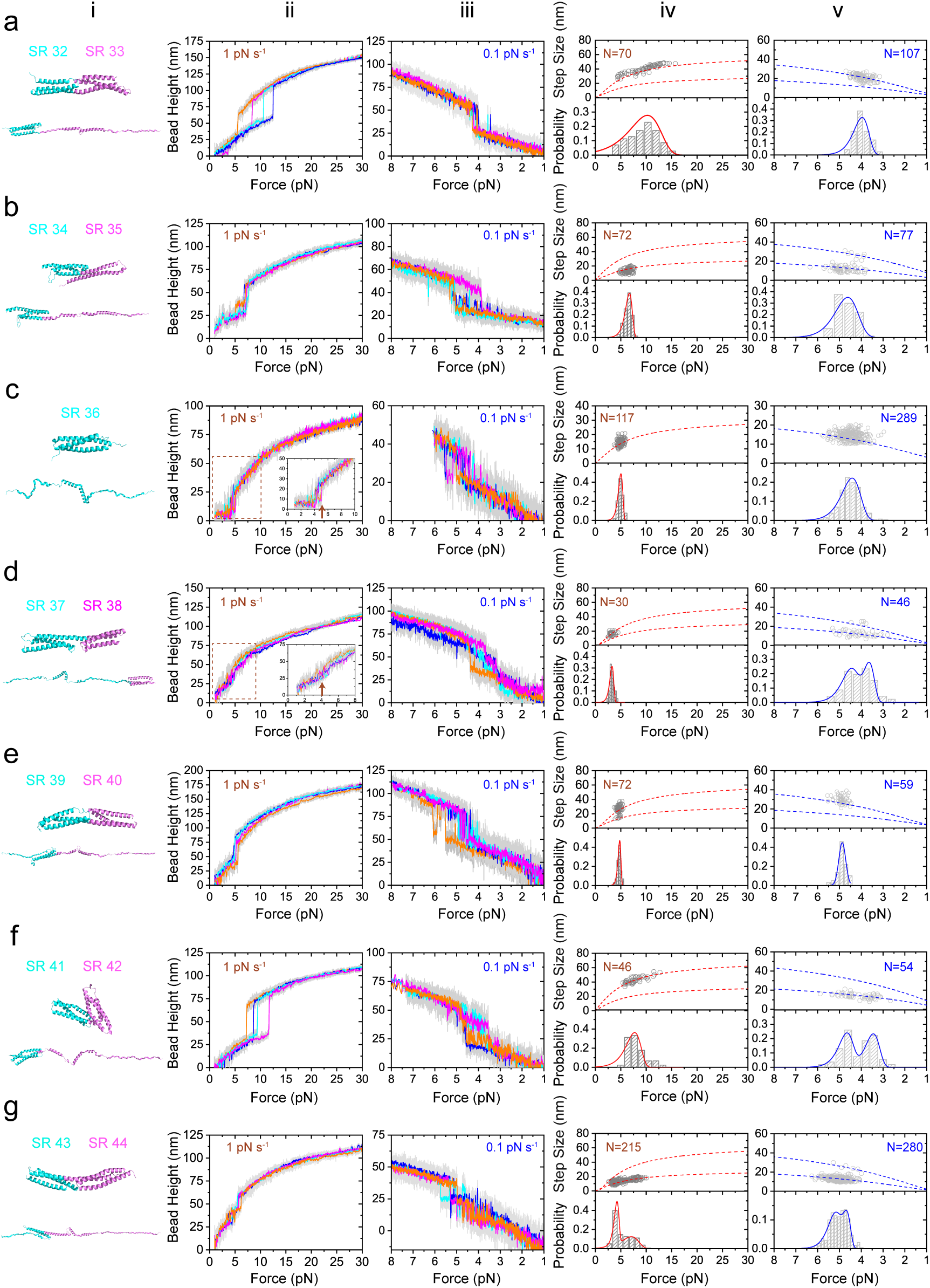
Force responses of giant nesprin-2 SR32-SR44 during linear force-loading. (**a-g**) Panel i: Representative snaps of corresponding SR domains before and after unfolding transition extracted from SMD simulations on AlphaFold2 predicted structures. Panels ii and iii: Representative force–bead height curves of the corresponding SR domains during linear force-increase (ii, 1 pN s^-1^) and force-decrease (iii, -0.1 pN s^-1^) scans. Raw data (gray) is 10-point FFT smoothed (colored lines). Panels iv and v: The resulting force–step size distributions of unfolding events (top, iv) and refolding events (top, v) obtained from the linear scans. The resulting normalized unfolding force (bottom, iv) and refolding force (bottom, v) distributions of the SRs. Number (*N*) of data points (represented by hollow circles) are indicated in the panels. The dashed colored lines are theoretical calculation of the force-dependent unfolding/refolding step sizes for corresponding SR domains. The colored solid lines are fitted force distributions based on Bell’s model (unfolding) or Arrhenius Law (refolding). For SR41-SR42, in the first scan, two SR unfolding events were observed. For the rest of cycles, only one SR unfolding/refolding event was observed and analyzed in panel iv and v.

**Fig. 5.**
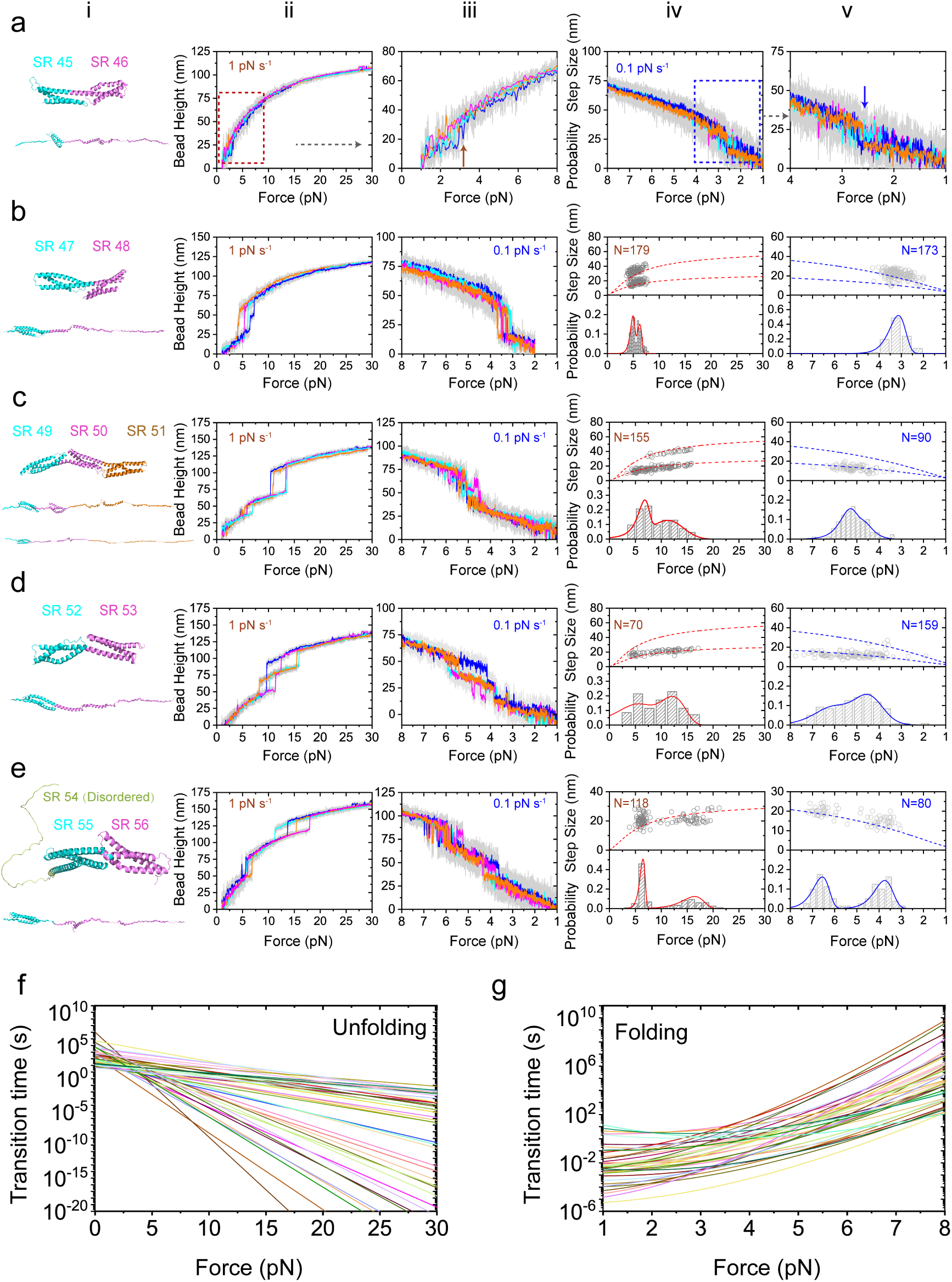
Force responses of giant nesprin-2 SR45-SR56 during linear force-loading. (**a**) For SR24-SR25, clear unfolding/refolding events was not observed. Panel iii and v are zoom-in of the force–bead height curves of ii and iv at low force regime, respectively. Arrows in iii and v indicate potential transition events. (**b-e**) Panel i: Representative snaps of corresponding SR domains before and after unfolding transition extracted from SMD simulations on AlphaFold2 predicted structures. Panels ii and iii: Representative force–bead height curves of the corresponding SR domains during linear force-increase (ii, 1 pN s^-1^) and force-decrease (iii, -0.1 pN s^-1^) scans. Raw data (gray) is 10-point FFT smoothed (colored lines). Panels iv and v: The resulting force–step size distributions of unfolding events (top, iv) and refolding events (top, v) obtained from the linear scans. The resulting normalized unfolding force (bottom, iv) and refolding force (bottom, v) distributions of the SRs. Number (*N*) of data points (represented by hollow circles) are indicated in the panels. The dashed colored lines are theoretical calculation of the force-dependent unfolding/refolding step sizes for corresponding SR domains. The colored solid lines are fitted force distributions based on Bell’s model (unfolding) or Arrhenius Law (refolding). (**f-g**) The extracted force-dependent unfolding (f) and refolding (g) lifetimes from experimentally obtained unfolding and refolding force distributions of the SR domains, based on Bell’s model and Arrhenius Law and analyses, respectively.

Figures 2-5 present characteristic force–bead height profiles of the giant nesprin-2 SR domains during linear force-increase (1 pN s^-1^) and force-decrease (-0.1 pN s^-1^) scans, as illustrated in Fig. 1d. Four representative domain curves are plotted (colored traces, with the corresponding raw data displayed in light gray following 10-point FFT smoothing, Figure 2-5 panels ii, iii). The data clearly demonstrate that all the SR domains in giant nesprin-2 unfold at forces below 25 pN. Subsequently, we analyzed both the force-dependent step sizes and the force distributions for unfolding and refolding transitions of the domains (Figs. 2-5, panels iv and v, respectively). The measured force-dependent unfolding and refolding step sizes exhibit excellent agreement with the theoretically calculated force-dependent step size values derived from the classical worm-like chain model for unfolded domains and the freely-jointed chain model for folded domains[51] (indicated by the dashed colored curves in panel iv and v, Fig. 2-5). The normalized distributions of unfolding and refolding forces for individual (or multiple) domains were well fitted based on Bell’s model and Arrhenius kinetics, respectively [52] (the solid colored curves in panel iv and v, Figs. 2-5), from which we can then extract the force-dependent unfolding and refolding lifetimes for each domain [4, 53] (Fig. 5f&g; see Methods: Arrhenius Law and Bell’s model analyses). Notably, in the absence of a resolved structure for giant nesprin-2, we calculated the rigid-body length (*b*_0_) of each domain using energy-minimized predicted structures from SMD simulations (Supplementary Figs. 2-6, see Methods: AlphaFold structural prediction, Steered molecular dynamics simulations, Theoretical force-dependent step sizes).

### Unstructured or mechanically weak SR domains within the giant nesprin-2

We directly stretched all 56 SRs and successfully detected clear force responses for 49 out of the 56 SRs, as shown in Figs. 2-5. For the remaining six SRs (SR20, SR24-SR25, SR45-SR46, and SR54), no clear single-molecule stretching signals were observed. Specific control signals, such as the SpyTag unzipping/rezipping dynamics at ∼30 pN, were observed in all of these constructs (Supplementary Fig. 1). We hypothesize that the lack of detectable protein unfolding signals in these constructs may result from extremely low mechanical stability, intrinsic disorder in the domains, potential segmentation, or protein aging effects.

For SR24-SR25 and SR45-SR46, by zooming in the fluctuations at low forces ≤2 pN, we observed some of the domain unfolding events at ∼2 pN or lower during linear force-increase scans (1 pN s*^−^*^1^), followed by refolding of the domains at ∼2 pN during linear force-decrease scans (-0.1 pN s*^−^*^1^). These observations indicate that these domains possess exceptionally low mechanical stability, causing them to readily unfold at forces around 1-2 pN. For SR20-SR21, only a single unfolding event was observed, corresponding to the unfolding of one SR domain. AlphaFold2 structural prediction and SMD simulation suggested that SR20-SR21 adopt stable folded conformations in the absence of force. These results collectively indicate that SR20 may also be mechanically weak and unfolds at forces ≤2 pN. Together, these results suggest that a subset of SR domains within the giant nesprin-2 protein is either unstructured or mechanically weak, likely existing in an unfolded state with small mechanical forces or thermal fluctuations.

In addition, to exclude potential segmentation effects, we analyzed longer protein constructs containing multiple neighboring SRs. For instance, the construct of SR9-SR19, in which we observed eleven unfolding events corresponding to SR9-SR19 (Supplementary Fig. 7a). The SR52-SR56 construct exhibited four unfolding events corresponding to SR52, SR53, SR55, and SR56 (Supplementary Fig. 7b), which is consistent with both the disordered prediction and the undetectable experimental observation of SR54 in the SR54-56 construct. The unfolding profiles of the longer constructs fully consisted of the segmented constructs. Here, we further excluded the possibility that some of the undetectable domains did not result from protein aging effects during experiments.

### Force-dependent lifetime of the giant nesprin-2 SRs

To gain further insight into the mechanical stability of giant nesprin-2, we extracted the force-dependent unfolding and refolding lifetimes from experimentally obtained unfolding and refolding force distributions [4, 53] (Fig. 5f&g; see Methods: Arrhenius Law and Bell’s model analyses). To ensure the accuracy of the extracted force-dependent transition lifetimes, we directly quantified the force-dependent lifetime of SR domains by implementing either a force-jump cycle procedure (Fig. 6a) or an equilibrium force measurement procedure (Fig. 6b-e).

**Fig. 6.**
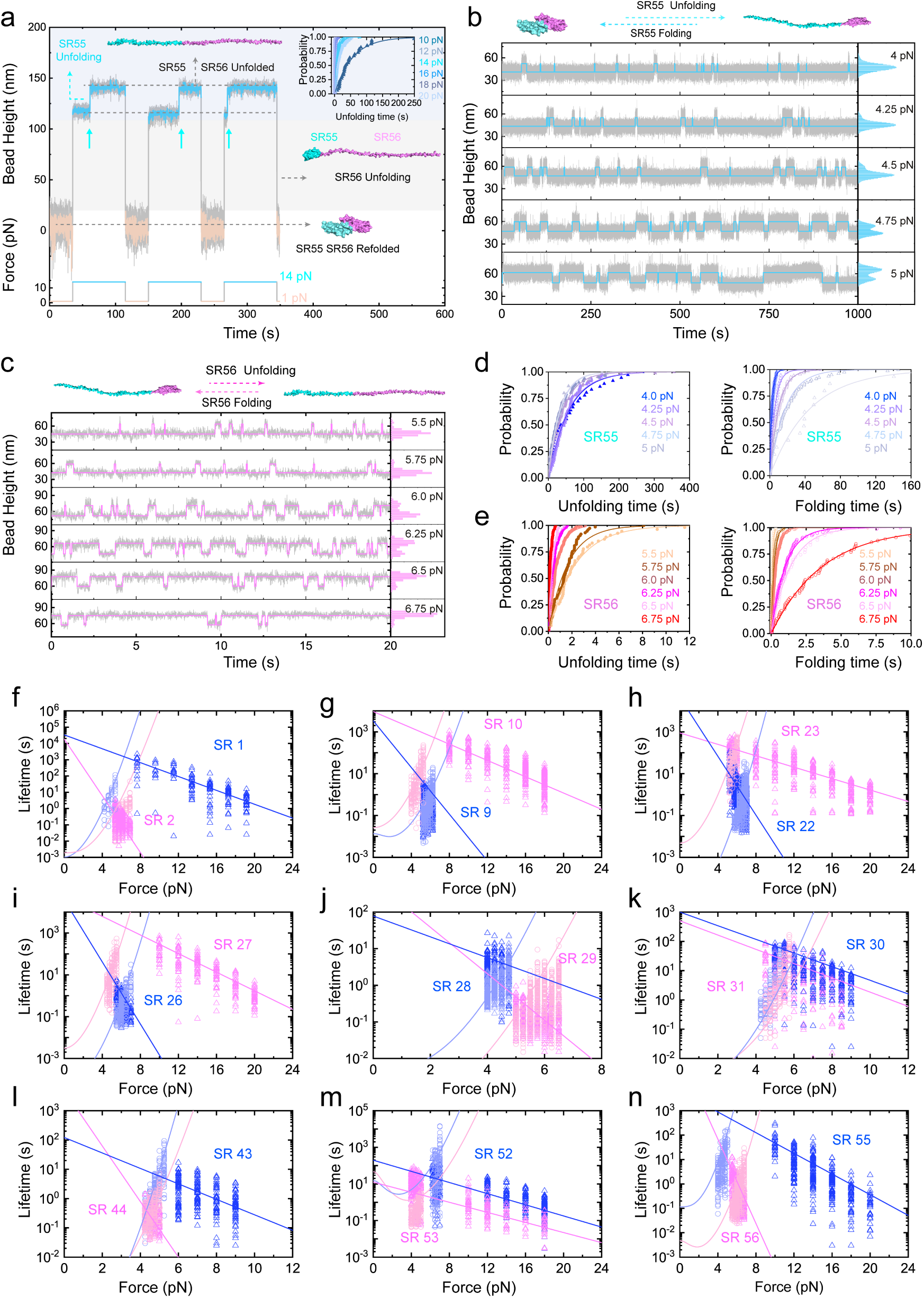
Force-dependent lifetimes of the giant nesprin-2. (**a**) Representative time–bead height curves of the SR54-SR56 during force-jump scans between 1 pN (orange) and 14 pN (cyan). The upward arrows indicate the unfolding events of SR55 at 14 pN. The SR56 unfolds during force-increase from 1 pN to 14 pN. The inset panel: The resulting unfolding time probability of the SR55 at forces of 10-20 pN. The lines are fitting curves (*p*(*t*) =) of the probability distribution. (**b-c**) Representative time– bead height curves of the SR54-SR56 during constant-force measurement at forces of 4-7 pN. Unfolding/refolding dyanmics of SR55 (b) and SR56 (c) at these forces are analyzed. (**d**) The resulting unfolding/refolding time probability of the SR55 (top panel) and SR56 (bottom panel) at forces of 4-7 pN. The lines are corresponding fitting curves of the probability distribution. The raw data (gray) is 10-point FFT smoothed (colored lines). (**f-n**) The resulting force-dependent unfolding and refolding lifetimes of the SR domains. Symbols indicate the experimentally direct obtained force-dependent lifetimes from the force-jump and constant force measurements. Lines indicate the extracted force-dependent lifetimes based on force-loading measurements on these SRs.

In each force-jump cycle, the domains were initially maintained at low forces (∼1 pN), to preserve their folded conformation. Subsequently, the force was abruptly elevated to a predetermined level and maintained until unfolding events were detected (e.g., SR55 unfolding at 14 pN in Fig. 6a). Following unfolding, the force was quickly decreased back to ∼1 pN, where it was held for approximately 30 seconds to facilitate domain refolding, recreating fully folded domains before the subsequent cycles. The dwell times (Δ*t*) prior to unfolding were systematically recorded for each cycle, and the domain lifetimes wrere determined by fitting the time-dependent unfolding probability based on the dwell time distribution using bootstrap analysis (e.g., SR55 unfolding under forces 10-20 pN, inset panel in Fig. 6a). Conversely, a reverse force-jump cycle, i.e., jumping from a high force where the domain was kept in unfolded state to a target lower force to monitor the refolding transition, could be employed to obtain the force-dependent refolding time of the SR domains (Supplementary Figs. 8-15).

In addition to the force-jump-cycle measurements, we also employed equilibrium-force procedures to quantify the unfolding and refolding lifetimes at equilibrium forces (Fig. 6b-e, Supplementary Figs. 8-15). In these procedures, the force was maintained at equilibrium values for sufficient durations to capture multiple dynamic unfolding and refolding transitions of the domains. The unfolding and refolding transition times at equilibrium forces were then derived by fitting the time-dependent unfolding and refolding probability to the dwell time distribution (e.g., SR55 and SR56 at 4-7 pN, Fig. 6b-e).

Figures 6f-n present the force-dependent unfolding and refolding lifetime obtained from 18 randomly selected SR domains out of the 56 in giant nesprin-2. Clearly, the directly measured force-dependent unfolding and refolding lifetimes (colored symbols) show excellent agreement with the extracted force-dependent unfolding and refolding lifetimes from force-loading experiments (colored lines). Based on this consistency, we subsequently employed the extracted force-dependent unfolding and refolding lifetimes for the analysis of the remaining SR domains. The unfolding and refolding lifetimes of the domains are distributed over a wide spectrum, resulting in distinct conformational dynamics of the domains within a few pN. This characteristic suggests the potential molecular shock absorber role of giant nesprin 2 (detailed in Fig. 7).

**Fig. 7.**
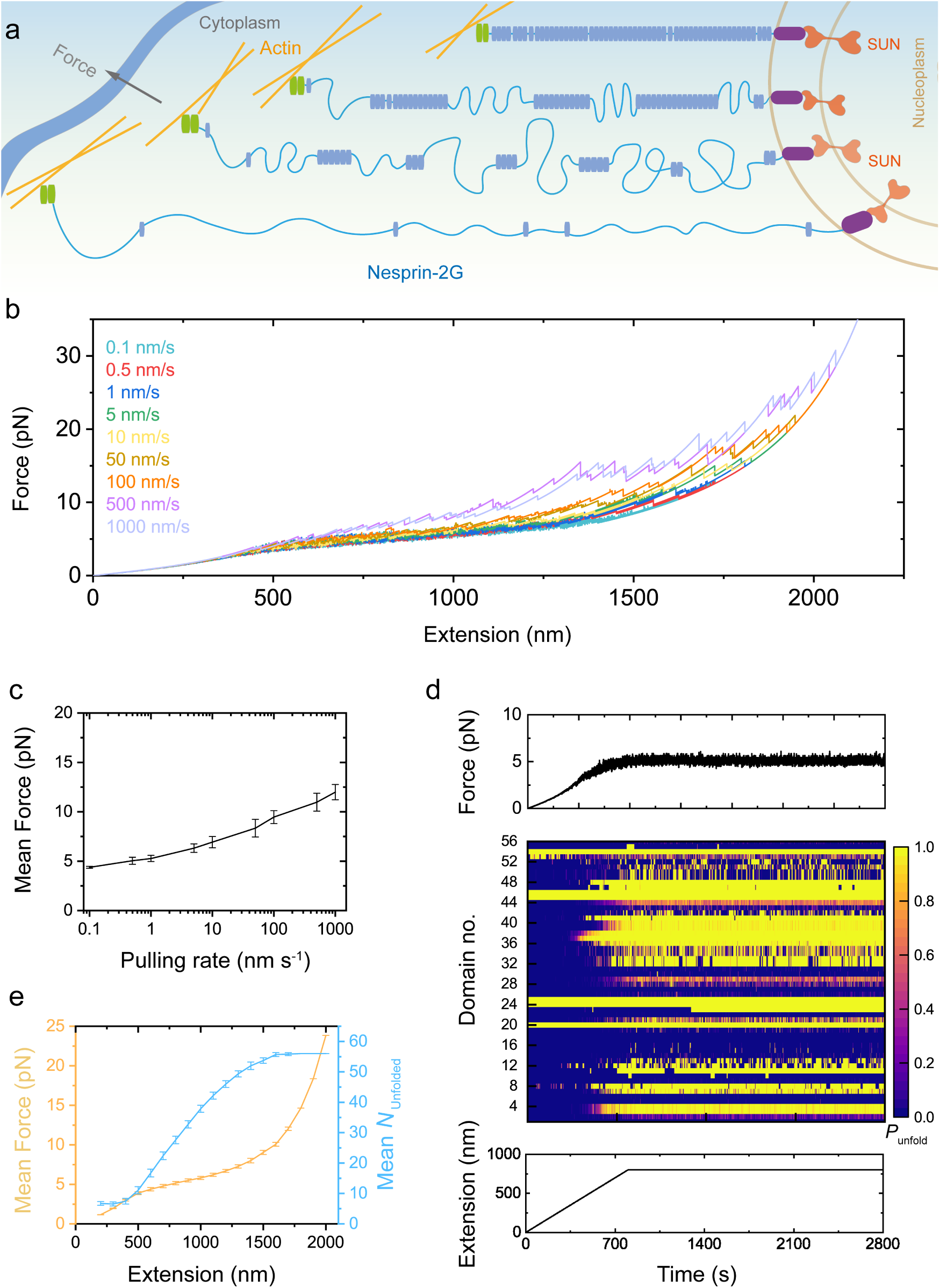
Giant nesprin-2 as a molecular shock absorber revealed by kinetic simulation. (**a**) Illustration of the giant nesprin-2 as a molecular shock absorber connecting the cytoskeleton and nucleoskeleton, by stochastic force-dependent unfolding and refolding dynamics of the numerous SR domains. (**b**) Representative extension–force curves of the full-length giant nesprin-2 during constant pulling with a wide range of pulling rates from 0.1 nm s^-1^ to 1000 nm s^-1^ in kinetic simulation. (**c**) The pulling rate dependent mean forces on the giant nesprin-2, obtained by kinetic simulation analysis. (**d**) An example force dynamics (top) and domain dynamics (middle) during the extension increasing and clamping processes. (**e**) The mean forces and and the number of unfold domains on giant nespirn-2 at different extensions of the nesprin-2.

### Asymmetric mechanical stability of the giant nesprin-2

As shown in the force-dependent unfolding step size curves and normalized unfolding force distributions (Fig. 2-5), the mechanical responses of giant nesprin-2’s SR domains can be classified into three distinct mechanical groups, besides those that are intrinsically disordered or extremely weak (Fig. 8, top panel). First, in the mechanically weak group, the domains primarily unfold at ∼ 5 pN. This group includes SR2-SR4, SR7-SR9, SR12-SR13, SR21-SR22, SR26, SR28-SR29, SR34-SR40, SR41-42, SR43-SR44, SR47-SR48, SR51, SR53, and SR56, accounting for 50% of the 56 SR domains. Considering the six intrinsically disordered or extremely weak domains, over 60% of the SR domains in giant nesprin-2 are mechanically weak and undergo rapid unfolding-refolding dynamics within forces of just a few pN. Second, approximately 25% of the domains (12 out of 56), specifically SR5-SR6, SR11, SR14-SR16, SR19, SR30-SR33, SR49-SR50, and SR52, mainly unfold at forces around ∼ 10 pN. Third, about 10% of the domains (7 out of 56) require forces exceeding ≥ 15 pN for unfolding. Interestingly, the mechanical weak domains (unfold rapidly at ≤5 pN) are mainly located at the C-terminal half, leading to an asymmetric mechanical stability profile in giant nesprin-2 (Fig.8, top panel).

**Fig. 8.**
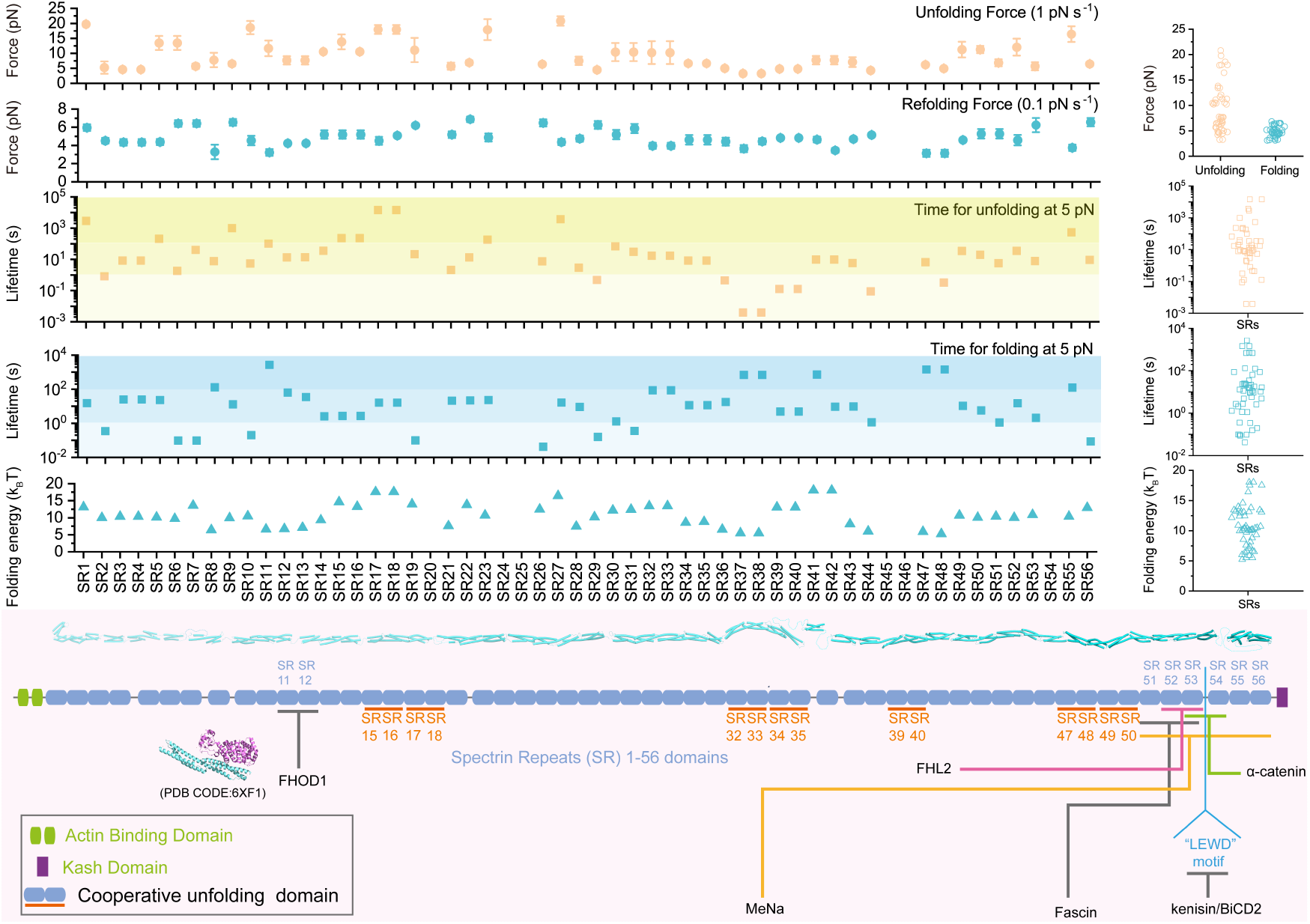
Mechanical stability of the giant nesprin-2 at a glance. First and second panels: The peak unfolding and refolding forces of all the SR domains of the giant nesprin-2, respectively. Third and fourth panels: The lifetimes of the folded SR (time taken for unfolding) and the unfolded SR (time taken for refolding) at a physiological force of 5 pN, respectively. Fifth panel: The zero-force folding energy of the SRs on the giant nesprin-2. Bottom panel: The illustration of the giant nesprin-2 protein and its known interacting partners.

### The thermostability of the domains of the giant nesprin-2

The force-dependent protein domain folding free energy can be calculated based on Boltzmann distribution:

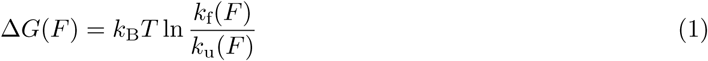

Then the zero-force folding free energy can be obtained:

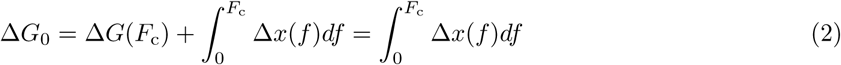

where the *F*_c_ is the critical force at which *k*_f_(*F*) = *k*_u_(*F*), the Δ*x*(*f*) is the force-dependent extension difference between the unfolded and folded states of a SR domain (i.e., the force-dependent step size). As summarized in Fig.8, the zero-force folding free energy of all 56 SR domains (except those mechanically extremely weak ones) were calculated.Importantly, while the force-dependent unfolding forces and unfolding rates reflect the mechanical stability of the domain, the zero-force folding free energy reflects the thermal stability of the domains. Most of the SRs are associated with a zero-force folding free energy of 5-20 *k*_B_*T* (i.e., 3-12 kcal mole^-1^), suggesting these domains are thermally stable in the absence of forces.

### The giant nesprin-2 as effective molecular shock absorbers

To estimate the tension in nesprin-mediated force transmission pathways connecting the cytoskeleton and nucleus (Fig. 7a), we performed stochastic kinetics simulations using an improved Gillespie algorithm[54]. This algorithm is based on the force-dependent unfolding and refolding rates of all 56 SR domains in giant nesprin-2 (Fig. 5f-g, see Methods: Kinetic simulation). Figure 7b displays the resulting extension-force curves of giant nesprin-2 at various pulling rates, covering the range of possible *in vivo* conditions. At each pulling rate, the curves exhibit a saw-tooth pattern. Each phase of increasing tension results from an extension increase without structural changes, while each abrupt force decrease or increase indicates the unfolding or refolding of an SR domain, which relaxes or increases the tension.

Importantly, across a wide range of pulling rates, from 0.1 nm s^-1^ to 10^3^ nm s^-1^, the average force in giant nesprin-2 remains around 5 pN over a large extension range (up to ∼1500 nm, which is about five times the length of nesprin in its folded state). Furthermore, at a constant extension ranging from 40 nm to 1500 nm, where nesprin-2 likely contains a mixture of folded and unfolded SR domains with diverse unfolding and refolding transition rates, the tension fluctuates around an average of 5 pN (Fig. 7). These results suggest that, through the stochastic unfolding and refolding of its force-bearing SR domains, giant nesprin-2 has the potential to function as a molecular mechanical shock absorber, maintaining force within a certain range along the entire force-transmission pathway it mediates during various cellular processes.

## Discussion

In this work, we combine direct single-molecule manipulation experiments with molecular dynamic simulations and AlphaFold structural predictions to investigate the giant nesprin-2 SR domains. Our results demonstrate that giant nesprin-2 undergoes dynamic unfolding and refolding transitions within physiological force ranges under force loading. This dynamic behavior of its numerous SR domains may function as a molecular shock absorber, effectively maintaining forces within the nesprin-2-mediated mechanotransduction pathway at approximately 5 pN, thereby modulating force transmission. Notably, unfolding and refolding of SR domains within a force range of a few pN suggests their versatile molecular regulation of the numerous SRs and their associated signaling partners during mechanosensing and mechanotransduction. This study establishes the molecular framework for understanding giant nesprin-2 mediated mechanotransduction.

The SR domain is a key structural feature of both the nesprin family and the spectrin superfamily, which includes the spectrin family, dystrophin/utrophin family, and *α*-actinin family, among others[3]. All of these proteins are mechanosensitive and physiologically subjected to mechanical forces *in vivo*. Therefore, understanding the response of SR domains to physiologically relevant forces–typically on the order of a few pN–is essential for uncovering their physiological functions. Among these SR superfamily proteins, giant nesprin-2, one of the largest SR-containing proteins, consists of 56 SR domains. Notably, over 60% of the SR domains in the giant nesprin-2 are mechanically weak domains that undergo rapid unfolding dynamics with forces ≤ 5 pN. The remaining of the SR domains rapidly unfold at forces within 20 pN during physiological force-loading. The differential mechanical stability of SR domains within physiological range enables giant nesprin-2 to dynamically adapt to varying mechanical microenvironments, thereby playing a crucial role in mechanotransduction.

Notably, 25% of the SR domains (14 out of 56, including SR15-SR16, SR17-SR18, SR32-SR33, SR34-SR35, SR39-SR40, SR47-SR48, and SR49-SR50) demonstrate pronounced cooperativity during mechanical unfolding. This is indicated by nearly simultaneous unfolding events, where two-domain unfolding events occur within less than 0.1 seconds during force-loading scans. Such cooperative unfolding behavior between adjacent SR domains likely represents a common feature of mechanosensitive proteinscontaining tandem SR domains [46]. The observed cooperativity may predominantly originate from salt-bridge networks mediated by inter-domain regions, which confer structural stability to both domains[55]. In support of this hypothesis, our structural analysis of the salt-bridge networks within giant nesprin-2, based on predicted structures, revealed that all cooperatively unfolding neighboring SR domains possess salt-bridge networks at their inter-domain linker regions (Supplementary Fig. 16). While our analysis focused on the most unambiguous cases of cooperativity (i.e., almost concurrent unfolding) in giant nesprin-2, inter-domain cooperativity effects may also likely present in some other domains, contributing to a certain degree of stabilization of the domains during mechanical unfolding. In addition the inter-domain salt bridges that mediating cooperativity among domains, other inter-domain interactions, such as *π* − *π* bond may also influence the mechanical stability and cooperativity of the SR domains. Notably, the *π* − *π* bond formed between SR41 and SR42 may enhance the SR41-SR42 cooperativity and mechanical stability (Supplementary Figure S17).

Nesprin family proteins are essential physical bridges between the cytoskeleton and the nucleus. Mechanical forces generated by cytoskeletal dynamics are transmitted to the nucleus via these bridges. Due to the stochastic unfolding and refolding of numerous force-bearing SR domains, giant nesprin-2 potentially serves as a molecular mechanical shock absorber, maintaining force levels within a certain range over physiological time scales. We propose that this force-buffering effect, driven by multiple-domain unfolding/refolding dynamics, may be conserved across various cellular mechanotransduction pathways, such as talin-mediated cell adhesion, dystrophin-mediated sarcolemma dynamics, and titin-mediated sarcomere dynamics[53, 56, 57]. Interestingly, all these multiple-domain unfolding/refolding dynamics result in average forces within a few pN, which aligns with the currently estimated forces acting on individual proteins within these mechanotransduction pathways[40, 58–72].

Nesprins function not only as critical mediators of mechanotransmission but also as essential signaling hubs coordinating mechanotransduction (Fig. 8, bottom panel). An expanding array of signaling partners has been reported in recent years[73–77, 77–80]. For instance, formin homology domain-containing protein 1 (FHOD1), directly binds the force-bearing spectrin repeat (SR) domains of nesprins via its N-terminal SR11-SR12 domains[73]. Disruption of this interface abolishes nuclear migration, underscoring its necessity for force transmission. Notably, FHOD1 also interacts with SR17-SR18 of giant nesprin-2[73]. Given that FHOD1 dimers anchor actin filament barbed ends, these multi-site interactions likely establish avidity-driven mechanoregulatory networks linking LINC complexes to cytoskeletal forces. A critical outstanding question is how physiological mechanical forces modulate FHOD1–giant nesprin-2 interactions. Elucidating this mechanism could provide critical insights into the architecture and adaptability of these mechanosensitive networks.

Interestingly, the SR49-SR56 domains at the C-terminal region of giant nesprin-2 function as a multi-functional signaling hub, playing critical roles in the cross-talk among microtubules, actin filaments, and the nucleoskeleton[74–77, 77–80]. This region contains a conserved W-acidic LEWD motif, which facilitates direct interaction with the light chain of kinesin-1[74–76, 80]. A recent in vitro study demonstrated that, together with MAP7 proteins, mini nesprin-2G directly links active kinesin-1 motors to F-actin, facilitating actin transport along microtubule tracks[74]. During neuronal migration, giant nesprin-2 recruits BicD2, a key dynein/kinesin adaptor, to the nuclear membrane via its C-terminal region. Disruption of the BicD2–giant nesprin-2 interaction significantly impairs nuclear movement[75, 76]. In epithelial–mesenchymal transitions (EMTs), the C-terminal region of giant nesprin-2 also recruits *α*-catenin to the nucleus, forming a supramolecular complex that includes *β*-catenin and emerin[77]. This recruitment attenuates Wnt/*β*-catenin signaling[77]. In patient-derived cutaneous squamous cell carcinoma cells, the actin regulatory protein Mena, a member of the Ena/VASP family, binds to giant nesprin-2 at this region on the nuclear membrane[78]. The Mena–giant nesprin-2 interaction regulates nuclear architecture, chromatin repositioning, and gene expression. Depletion of Mena leads to dissociation of nesprin-2 from actin and lamin A/C[78]. In cardiomyocytes, both telethonin and four-and-a-half LIM domain (FHL)-2, key regulators of sarcomere dynamics, bind to the C-terminal region of giant nesprin-2[79]. Mutations in nesprin-2, telethonin, and FHL-2, identified in patients with EDMD, DCM, and hypertrophic cardiomyopathy, impair these interactions[79]. Together, these diverse interactions of the C-terminal SR domains of giant nesprin-2 underscore the need for future studies on the molecular mechanisms underlying the mechano-chemical regulation of these critical interactions (Fig.8).

## Methods

### Plasmids and proteins

28 expression plasmids were engineered for single-molecule stretching experiments. The insert SR domain genes were amplified by PCR from template DNA using Q5 High-Fidelity DNA Polymerase (NEB) or by gene synthesis (Sangon). Then the DNA fragments were cloned into a pET151 or a pET28a vector using seamless assembly and confirmed by Sanger sequencing. For protein production, each plasmid was co-transformed with a BirA biotin ligase expression vector into *Escherichia coli* BL21 (DE3) cells. Cultures were grown in LB-medium supplemented with D-Biotin, and recombinant proteins were purified by Ni-NTA affinity chromatography via their N-terminal 6His-tags. The detailed sequences of the corresponding protein constructs are provided in Supplementary Note 1. Detailed protocols for protein expression and purification steps can be found in Supplementary Note 2. The names of the plasmid constructs and domains included within the constructs with the corresponding residues based on uniProt database (Q6ZWQ0-1) are listed below.

1. pET151-Avi-(SR1-SR2)_Nesprin-2G_-Spy, 299 − 474;
2. pET151-Avi-(SR3-SR4)_Nesprin2G_-Spy, 475 − 680;
3. pET151-Avi-(SR5-SR7)_Nesprin2G_-Spy, 727 − 1030;
4. pET151-Avi-(SR8)_Nesprin2G_-Spy, 1062 − 1211;
5. pET151-Avi-(SR9-SR10)_Nesprin2G_-Spy, 1262 − 1409;
6. pET151-Avi-(SR11-SR13)_Nesprin2G_-Spy, 1410 − 1728;
7. pET151-Avi-(SR14-SR16)_Nesprin2G_-Spy, 1729 − 2026;
8. pET151-Avi-(SR17-SR19)_Nesprin2G_-Spy, 2027 − 2350;
9. pET151-Avi-(SR20-SR21)_Nesprin2G_-Spy, 2422 − 2610;
10. pET151-Avi-(SR22-SR23)_Nesprin2G_-Spy, 2611 − 2821;
11. pET151-Avi-(SR24-SR25)_Nesprin2G_-Spy, 2822 − 3027;
12. pET151-Avi-(SR26-SR27)_Nesprin2G_-Spy, 3028 − 3239;
13. pET151-Avi-(SR28-SR29)_Nesprin2G_-Spy, 3240 − 3456;n
14. pET151-Avi-(SR30-SR31)_Nesprin2G_-Spy, 3457 − 3669;
15. pET151-Avi-(SR32-SR33)_Nesprin2G_-Spy, 3670 − 3870;
16. pET151-Avi-(SR34-SR35)_Nesprin2G_-Spy, 3871 − 4074;
17. pET151-Avi-(SR36)_Nesprin2G_-Spy, 4218 − 4337;
18. pET151-Avi-(SR37-SR38)_Nesprin2G_-Spy, 4507 − 4714;
19. pET151-Avi-(SR39-SR40)_Nesprin2G_-Spy, 4715 − 4929;
20. pET151-Avi-(SR41-SR42)_Nesprin2G_-Spy, 4930 − 5150;
21. pET151-Avi-(SR43-SR44)_Nesprin2G_-Spy, 5151 − 5377;
22. pET151-Avi-(SR45-SR46)_Nesprin2G_-Spy, 5378 − 5576;
23. pET151-Avi-(SR47-SR48)_Nesprin2G_-Spy, 5577 − 5786;
24. pET151-Avi-(SR49-SR51)_Nesprin2G_-Spy, 5787 − 6122;
25. pET151-Avi-(SR52-SR53)_Nesprin2G_-Spy, 6110 − 6362;
26. pET151-Avi-(SR54-SR56)_Nesprin2G_-Spy, 6433 − 6822;
27. pET151-Avi-(SR52-SR56)_Nesprin2G_-Spy, 6110 − 6822;
28. pET151-Avi-(SR09-SR19)_Nesprin2G_-Spy, 1262 − 2350;

### Magnetic tweezers manipulation

A vertical magnetic tweezers setup was integrated with a disturbance-free, rapid solution-exchange flow channel to perform *in vitro* protein stretching experiments [47, 48, 81–84]. All experiments were conducted in a solution containing 1X PBS, 1% BSA, 2 mM DTT, 10 mM sodium L-ascorbate at 22 ± 1 ^o^C. The force calibration of the magnetic tweezers exhibits a 10% uncertainty, primarily arising from heterogeneity in the diameters of paramagnetic beads [47]. Bead height determination introduces an additional uncertainty of ∼ 2-5 nm due to thermal fluctuations of the tethered bead and the limited resolution of the imaging system [81].

### Theoretical analysis of force-dependent step sizes

The force-extension behavior of a folded domain is dictated by the rotational fluctuations of a characteristic rigid body with a length *b*_0_. This parameter, *b*_0_, represents the separation between the two force application points and can be inferred from the PDB structure of the folded domain (Supplementary Fig. 2-6). The extension response of such a rigid body follows the freely-jointed chain (FJC) polymer model with a single segment, mathematically expressed as:

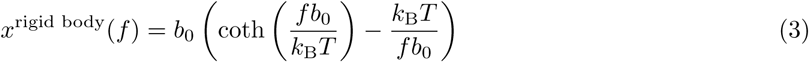

In contrast, the force-extension relationship of the unfolded domain is governed by the mechanical response of a flexible polypeptide chain. This behavior can be accurately described using the worm-like chain (WLC) polymer model, formulated by Marko and Siggia [51], with an intrinsic bending persistence length of approximately *A* ∼ 0.8 nm:

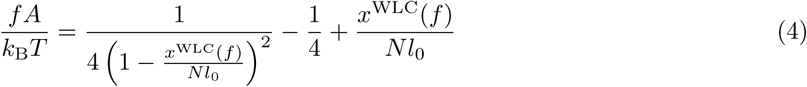

where *l*_0_ = 0.38 nm represents the contour length per amino acid residue, and *N* denotes the total number of residues in the unfolded domain.

As a result, the force-dependent step size associated with the unfolding transition corresponds to the differential extension of the domain before and after unfolding at the transition force, given by:

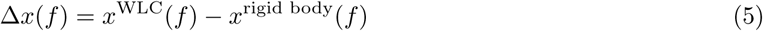

### Steered molecular dynamics simulations

The force field CHARMM36m[85] and water models TIP3P[86] and Gromacs 5.1.4 software[87] were selected for steered molecular dynamics (SMD) simulations. The native structures of nesprin-2 SR domain are based on the predictions of Alphafold2 (Supplementary Fig. 2-6). The system was solved in a 150 mM NaCl solution while ensuring charge equilibrium. Initial energy minimization was performed using the steepest descent method. Subsequently, 1 ns of NPT (constant Number of particles, Pressure, and Temperature) system equilibration was conducted, harmonic position restriction was applied to the heavy atoms of the protein during this equilibration process. The temperature was held at 298 K while the pressure was kept at 1 bar. A harmonic spring with a spring constant of *k* = 100 kJ mol^-1^·nm is used to generate tension between the N and C termini of SR domains. The N-C distance was recorded while the spring is moving at a constant speed *v* = 1 nm ns^-1^[88]. The time evolutions of the domains in SMD are included in Supplementary Mov. 1-26.

To investigate the structural basis of the mechanical responses observed in SMD simulations. Potential salt bridges were estimated based on the distance (typically *<* 4 °A) between positively charged residues (ARG, HIS, LYS) and a negatively charged residues (ASP, GLU). Before salt-bridge analysis, each SR structure was fully relaxed by energy minimization with Gromacs.

### Arrhenius Law and Bell’s model analyses

The kinetic parameters of the SR domains were encoded in the unfolding and refolding force distributions observed under linear force-loading. Accordingly, we extracted these parameters by fitting the normalized force distributions. At a given loading rate *r* (positive for force increase and negative for force decrease), the unfolding or refolding distribution *ρ*(*f*) is given by:

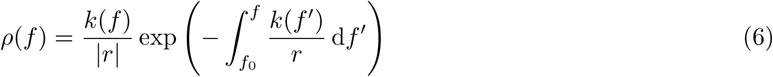

where *k*(*f*) denotes the force-dependent unfolding or refolding rate and *f*_0_ represents the force at which the transition probability is zero (e.g., 0 pN for unfolding and several tens of pN for refolding).

Here, we assume that the unfolding rate *k*_u_(*f*) of the SR domains follows the Bell’s model: 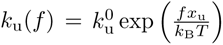, where *k_f_*^0^ is the zero-force unfolding rate and *x*_u_ is the transition distance between the native state and the transition state. In contrast, the refolding rate is modeled using the Arrhenius law: 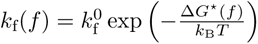, where *k*_u_^0^ is the zero-force refolding rate and Δ*G*^*^(*f*) is the free energy difference between the transition state and the unfolded state. In detail, 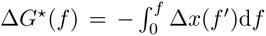 and the force-dependent transitions distance Δ*x*(*f*) = *x*^*^(*f*) − *x*_unfolded_(*f*). To evaluate these distances, the transition state is treated as a rigid core of length *b*^*^, for which the extension *x*^*^(*f*) is described by the freely-jointed chain model, while the unfolded state is treated as a flexible peptide and *x*_unfolded_(*f*) can be calculated using the worm like chain model. The force dependent transition time *τ* (*f*) = 1*/k*(*f*).

### Kinetic simulation analysis

Briefly, the 56 SR domains are treated as a one-dimensional lattice model. At a given extension of the giant nesprin-2, the force in the SRs is estimated based on the structural states (i.e., folded or unfolded) of the domains and the force-extension curves of the domains in the corresponding states. In addition, the intrinsically disordered domains and other linker regions are modeled as flexible chains which follow the WLC model [51]. In response to the force, each SR domain undergoes stochastic structural transitions based on its force dependent unfolding and refolding transition rates (Fig. 5f-g, Supplementary Table 1-2). Both the time to the next transition event and the domain involved in that transition are stochastically determined based on transition rates of each domain. After each transition, the structural states of the lattice were updated, and the resulting force was recalculated. Iteration of this process results in evolution of the structural states, causing force fluctuation in the giant nesprin-2, at a given extension. In addition, the same simulation performed on a time varying extension of the giant nesprin-2 can predict changes of tension and structural states of the protein domains at different pulling rates (i.e., the rate of total extension change).

## Author Contributions

S.L. and M.Y. conceived the study and designed the experiments. F.S., Y.Z. J.Y. and Z.Z. performed the single molecule experiments; Y.Z. performed the molecular dynamics simulations; J.Y. performed the kinetic simulations; F.S., S.L. and M.Y. prepared the plasmids and proteins; S.L., M.Y., H.C., S.F., Y.Z. and J.Y. analyzed and interpreted the data; S.L. and M.Y. wrote the paper with inputs from all authors.

## Acknowledgment

The research is funded by the National Natural Science Foundation of China (NSFC Grant NO. 12474202 and 32301094 to M.Y., NSFC Grant NO. 32271367 and 12204389 to S.L., NSFC Grant NO. 12474200 to H.C.) and the 111 Project (B16029). The authors acknowledge the insightful discussions with Prof. Jie Yan (National University of Singapore), Prof. Chenxu Wu (Xiamen University), Prof. Wei Chen and Prof. Yiting Zhou (Zhejiang University).

## Data availability

Unless otherwise stated, all data supporting the results of this study can be found in the article, Supplementary, and source data files. Sequence information of key constructs is included in Supplementary Information. AlphaFold2-predicted structures are available as source data files. The predicted models are subject to the AlphaFold 2 Server (EMBL-EBI) Terms (available at https://alphafold.ebi.ac.uk/terms). All structures were used strictly for non-commercial, scientific research purposes. Key plasmids are available upon request from the corresponding authors. Source data are provided in this paper.

## References

[1] Fletcher, D. A. & Mullins, R. D. Cell mechanics and the cytoskeleton. Nature 463, 485–492 (2010).

[2] Blanchoin, L., Boujemaa-Paterski, R., Sykes, C. & Plastino, J. Actin dynamics, architecture, and mechanics in cell motility. Physiological Reviews 94, 235–263 (2014).

[3] Liem, R. K. Cytoskeletal integrators: The spectrin superfamily. Cold Spring Harbor Perspectives in Biology 8 (2016).

[4] Le, S., Yu, M. & Yan, J. Mechanical regulation of tension-transmission supramolecular linkages. Current Opinion in Solid State and Materials Science 25, 100895 (2021).

[5] Maurer, M. & Lammerding, J. The driving force: Nuclear mechanotransduction in cellular function, fate, and disease. Annual Review of Biomedical Engineering 21, 443–468 (2019).

[6] Lee, Y. L. & Burke, B. LINC complexes and nuclear positioning. Seminars in Cell & Developmental Biology 82, 67–76 (2018).

[7] Burke, B. LINC complexes as regulators of meiosis. Current Opinion in Cell Biology 52, 22–29 (2018).

[8] Horn, H. F. et al. The linc complex is essential for hearing. The Journal of clinical investigation 123, 740–750 (2013).

[9] Sosa, B. A., Rothballer, A., Kutay, U. & Schwartz, T. U. Linc complexes form by binding of three kash peptides to domain interfaces of trimeric sun proteins. Cell 149, 1035–1047 (2012).

[10] Zhou, Z. et al. Structure of sad1-unc84 homology (sun) domain defines features of molecular bridge in nuclear envelope. The Journal of biological chemistry 287, 5317–5326 (2012).

[11] Crisp, M. et al. Coupling of the nucleus and cytoplasm: role of the linc complex. The Journal of cell biology 172, 41–53 (2006).

[12] Starr, D. A. & Han, M. Role of anc-1 in tethering nuclei to the actin cytoskeleton. Science 298, 406–409 (2002).

[13] Grady, R. M., Starr, D. A., Ackerman, G. L., Sanes, J. R. & Han, M. Syne proteins anchor muscle nuclei at the neuromuscular junction. Proceedings of the National Academy of Sciences of the United States of America 102, 4359–4364 (2005).

[14] Neumann, S. et al. Nesprin-2 interacts with *α*-catenin and regulates wnt signaling at the nuclear envelope. Journal of Biological Chemistry 285, 34932–34938 (2010).

[15] Chikashige, Y. et al. Meiotic proteins bqt1 and bqt2 tether telomeres to form the bouquet arrangement of chromosomes. Cell 125, 59–69 (2006).

[16] Swartz, R. K., Rodriguez, E. C. & King, M. C. A role for nuclear envelope-bridging complexes in homology-directed repair. Molecular biology of the cell 25, 2461–2471 (2014).

[17] Alam, S. G. et al. The mammalian linc complex regulates genome transcriptional responses to substrate rigidity. Scientific reports 6, 38063 (2016).

[18] Wang, S. et al. Mechanotransduction via the linc complex regulates dna replication in myonuclei. The Journal of cell biology 217, 2005–2018 (2018).

[19] Driscoll, T. P., Cosgrove, B. D., Heo, S.-J., Shurden, Z. E. & Mauck, R. L. Cytoskeletal to nuclear strain transfer regulates yap signaling in mesenchymal stem cells. Biophysical journal 108, 2783–2793 (2015).

[20] Elosegui-Artola, A. et al. Force triggers yap nuclear entry by regulating transport across nuclear pores. Cell 171, 1397–1410.e14 (2017).

[21] Uzer, G. et al. Sun-mediated mechanical linc between nucleus and cytoskeleton regulates catenin nuclear access. Journal of biomechanics 74, 32–40 (2018).

[22] Tajik, A. et al. Transcription upregulation via force-induced direct stretching of chromatin. Nature materials 15, 1287–1296 (2016).

[23] Zhang, Q. et al. Nesprin-1 and -2 are involved in the pathogenesis of emery dreifuss muscular dystrophy and are critical for nuclear envelope integrity. Human molecular genetics 16, 2816–2833 (2007).

[24] Gros-Louis, F. et al. Mutations in syne1 lead to a newly discovered form of autosomal recessive cerebellar ataxia. Nature genetics 39, 80–85 (2007).

[25] Attali, R. et al. Mutation of syne-1, encoding an essential component of the nuclear lamina, is responsible for autosomal recessive arthrogryposis. Human molecular genetics 18, 3462–3469 (2009).

[26] Puckelwartz, M. J. et al. Nesprin-1 mutations in human and murine cardiomyopathy. Journal of molecular and cellular cardiology 48, 600–608 (2010).

[27] Green, E. K. et al. Association at syne1 in both bipolar disorder and recurrent major depression. Molecular psychiatry 18, 614–617 (2013).

[28] Gerace, L. & Huber, M. D. Nuclear lamina at the crossroads of the cytoplasm and nucleus. Journal of structural biology 177, 24–31 (2012).

[29] Haque, F. et al. Sun1 interacts with nuclear lamin a and cytoplasmic nesprins to provide a physical connection between the nuclear lamina and the cytoskeleton. Molecular and cellular biology 26, 3738– 3751 (2006).

[30] Sjöblom, B., Salmazo, A. & Djinović-Carugo, K. *α*-actinin structure and regulation. Cellular and Molecular Life Sciences 65, 2688–2701 (2008).

[31] Djinović-Carugo, K., Young, P., Gautel, M. & Saraste, M. Structure of the alpha-actinin rod: molecular basis for cross-linking of actin filaments. Cell 98, 537–546 (1999).

[32] Zhang, Q. et al. Nesprins: a novel family of spectrin-repeat-containing proteins that localize to the nuclear membrane in multiple tissues. Journal of Cell Science 114, 4485–4498 (2001).

[33] Zhang, Q., Ragnauth, C., Greener, M. J., Shanahan, C. M. & Roberts, R. G. The nesprins are giant actin-binding proteins, orthologous to drosophila melanogaster muscle protein msp-300. Genomics 80, 473–481 (2002).

[34] Roux, K. J. et al. Nesprin 4 is an outer nuclear membrane protein that can induce kinesin-mediated cell polarization. Proceedings of the National Academy of Sciences of the United States of America 106, 2194–2199 (2009).

[35] Wilhelmsen, K. et al. Nesprin-3, a novel outer nuclear membrane protein, associates with the cytoskeletal linker protein plectin. The Journal of cell biology 171, 799–810 (2005).

[36] Rajgor, D. & Shanahan, C. M. Nesprins: from the nuclear envelope and beyond. Expert Reviews in Molecular Medicine 15 (2013).

[37] Arsenovic, P. T. et al. Nesprin-2g, a component of the nuclear LINC complex, is subject to myosin-dependent tension. Biophysical Journal 110, 34–43 (2016).

[38] Guilluy, C. et al. Isolated nuclei adapt to force and reveal a mechanotransduction pathway in the nucleus. Nature Cell Biology 16, 376–381 (2014).

[39] Déjardin, T., et al. Nesprins are mechanotransducers that discriminate epithelial–mesenchymal transition programs. Journal of Cell Biology 219 (2020).

[40] Ren, Y. et al. Genetically encoded mechano-sensors with versatile readouts and compact size. bioRxiv (2025).

[41] Denais, C. M. et al. Nuclear envelope rupture and repair during cancer cell migration. Science 352, 353–358 (2016).

[42] Davidson, P. M. et al. Nesprin-2 accumulates at the front of the nucleus during confined cell migration. EMBO reports 21 (2020).

[43] Jumper, J. et al. Highly accurate protein structure prediction with alphafold. Nature 596, 583–589 (2021).

[44] Zakeri, B. et al. Peptide tag forming a rapid covalent bond to a protein, through engineering a bacterial adhesin. Proceedings of the National Academy of Sciences 109 (2012).

[45] Le, S. et al. Disturbance-free rapid solution exchange for magnetic tweezers single-molecule studies. Nucleic Acids Research 43, e113–e113 (2015).

[46] Le, S. et al. Mechanotransmission and mechanosensing of human alpha-actinin 1. Cell Reports 21, 2714–2723 (2017).

[47] Chen, H. et al. Improved high-force magnetic tweezers for stretching and refolding of proteins and short DNA. Biophysical Journal 100, 517–523 (2011).

[48] Le, S., Liu, R., Lim, C. T. & Yan, J. Uncovering mechanosensing mechanisms at the single protein level using magnetic tweezers. Methods 94, 13–18 (2016).

[49] Jo, M. H. et al. Determination of single-molecule loading rate during mechanotransduction in cell adhesion. Science 383, 1374–1379 (2024).

[50] Hu, Y. et al. Dna-based forcechrono probes for deciphering single-molecule force dynamics in living cells. Cell 187, 3445–3459.e15 (2024).

[51] Marko, J. F. & Siggia, E. D. Stretching dna. Macromolecules 28, 8759–8770 (1995).

[52] Bell, G. I. Models for the specific adhesion of cells to cells. Science 200, 618–627 (1978).

[53] Le, S. et al. Dystrophin as a molecular shock absorber. ACS Nano 12, 12140–12148 (2018).

[54] Gillespie, D. T. A general method for numerically simulating the stochastic time evolution of coupled chemical reactions. Journal of Computational Physics 22, 403–434 (1976).

[55] Le, S. et al. Single-molecule force spectroscopy reveals intra- and intermolecular interactions of Caenorhabditis elegans HMP-1 during mechanotransduction. Proceedings of the National Academy of Sciences of the United States of America 121, e2400654121 (2024).

[56] Yao, M. et al. The mechanical response of talin. Nature Communications 7, 11966 (2016).

[57] Yuan, G. et al. Elasticity of the transition state leading to an unexpected mechanical stabilization of titin immunoglobulin domains. Angewandte Chemie International Edition 56, 5490–5493 (2017).

[58] Grashoff, C. et al. Measuring mechanical tension across vinculin reveals regulation of focal adhesion dynamics. Nature 466, 263–266 (2010).

[59] Borghi, N. et al. E-cadherin is under constitutive actomyosin-generated tension that is increased at cell-cell contacts upon externally applied stretch. Proceedings of the National Academy of Sciences of the United States of America 109, 12568–73 (2012).

[60] Morimatsu, M., Mekhdjian, A. H., Adhikari, A. S. & Dunn, A. R. Molecular tension sensors report forces generated by single integrin molecules in living cells. Nano Letters 13, 3985–3989 (2013).

[61] Wang, X. & Ha, T. Defining single molecular forces required to activate integrin and notch signaling. Science 340, 991–994 (2013).

[62] Zhang, Y., Ge, C., Zhu, C. & Salaita, K. DNA-based digital tension probes reveal integrin forces during early cell adhesion. Nature Communications 55167 (2014).

[63] Blakely, B. L. et al. A dna-based molecular probe for optically reporting cellular traction forces. Nature Methods 11, 1229–1232 (2014).

[64] Austen, K. et al. Extracellular rigidity sensing by talin isoform-specific mechanical linkages. Nature Cell Biology 17, 1597–1606 (2015).

[65] Ringer, P. et al. Multiplexing molecular tension sensors reveals piconewton force gradient across talin-1. Nature Methods 14, 1090–1096 (2017).

[66] Chang, A. C. et al. Single molecule force measurements in living cells reveal a minimally tensioned integrin state. ACS Nano 10, 10745–10752 (2016).

[67] Cai, D. et al. Mechanical feedback through e-cadherin promotes direction sensing during collective cell migration. Cell 157, 1146–1159 (2014).

[68] Price, A. J. et al. Mechanical loading of desmosomes depends on the magnitude and orientation of external stress. Nature Communications 9, 5284 (2018).

[69] Li, H. et al. A reversible shearing dna probe for visualizing mechanically strong receptors in living cells. Nature Cell Biology 23, 642–651 (2021).

[70] Wang, W. et al. Hydrogel-based molecular tension fluorescence microscopy for investigating receptor-mediated rigidity sensing. Nature Methods 20, 1780–1789 (2023).

[71] Brockman, J. M. et al. Live-cell super-resolved paint imaging of piconewton cellular traction forces. Nature Methods 17, 1018–1024 (2020).

[72] Kubow, K. E. et al. Mechanical forces regulate the interactions of fibronectin and collagen i in extracellular matrix. Nature Communications 6, 8026 (2015).

[73] Lim, S. M., Cruz, V. E., Antoku, S., Gundersen, G. G. & Schwartz, T. U. Structures of fhod1-nesprin1/2 complexes reveal alternate binding modes for the fh3 domain of formins. Structure 29, 540–552.e5 (2021).

[74] Sahabandu, N., et al. Active microtubule-actin crosstalk mediated by a nesprin-2g-kinesin complex. bioRxiv (2024).

[75] Gonçalves, J. C., Quintremil, S., Yi, J. & Vallee, R. B. Nesprin-2 recruitment of bicd2 to the nuclear envelope controls dynein/kinesin-mediated neuronal migration in vivo. Current Biology 30, 3116– 3129.e4 (2020).

[76] Yi, J. et al. Role of nesprin-2 and ranbp2 in bicd2-associated brain developmental disorders. PLOS Genetics 19, e1010642 (2023).

[77] Gottardi, C. J. & Luxton, G. G. Nesprin-2g tension fine-tunes wnt/*β*-catenin signaling. Journal of Cell Biology 219,e202009042 (2020).

[78] Li Mow Chee, F., et al. Mena regulates nesprin-2 to control actin–nuclear lamina associations, trans-nuclear membrane signalling and gene expression. Nature Communications 14, (2023).

[79] Li, C. et al. Nesprin-2 is a novel scaffold protein for telethonin and fhl-2 in the cardiomyocyte sarcomere. Journal of Biological Chemistry 300, 107254 (2024).

[80] Zhou, C., Wu, Y. K., Ishidate, F., Fujiwara, T. K. & Kengaku, M. Nesprin-2 coordinates opposing microtubule motors during nuclear migration in neurons. Journal of Cell Biology 223,e202405032 (2024).

[81] Le, S., Yu, M. & Yan, J. Phosphorylation reduces the mechanical stability of the *α*-catenin/*β*-catenin complex. Angewandte Chemie International Edition 58, 18663–18669 (2019).

[82] Zhang, Y. et al. Multi-domain interaction mediated strength-building in human *α*-actinin dimers unveiled by direct single-molecule quantification. Nature Communications 15, 6151 (2024).

[83] Wang, Y. et al. Mechanically weak and highly dynamic state of mechanosensitive titin ig domains induced by proline isomerization. Nature Communications 16, 2771 (2025).

[84] Zhang, Y. et al. Anomalous force-dependent transition rates unveil dual pathways in folding and unfolding dynamics of acyl-coenzyme a binding protein. The Journal of Physical Chemistry Letters 16, 2479–2486 (2025).

[85] Huang, J. et al. Charmm36m: an improved force field for folded and intrinsically disordered proteins. Nature methods 14, 71–73 (2017).

[86] Jorgensen, W. L. Quantum and statistical mechanical studies of liquids. 10. transferable intermolecular potential functions for water, alcohols, and ethers. application to liquid water. Journal of the American Chemical Society 103, 335–340 (1981).

[87] Van Der Spoel, D. et al. Gromacs: fast, flexible, and free. Journal of computational chemistry 26, 1701–1718 (2005).

[88] Fersht, A. R. & Daggett, V. Protein folding and unfolding at atomic resolution. Cell 108, 573–582 (2002).

