## Supplementary Information File for "Mechanical Responses of the Giant Nesprin-2"

### Supplementary Information for
### Mechanical Responses of the Giant Nesprin-2

(Dated: May 30, 2025)

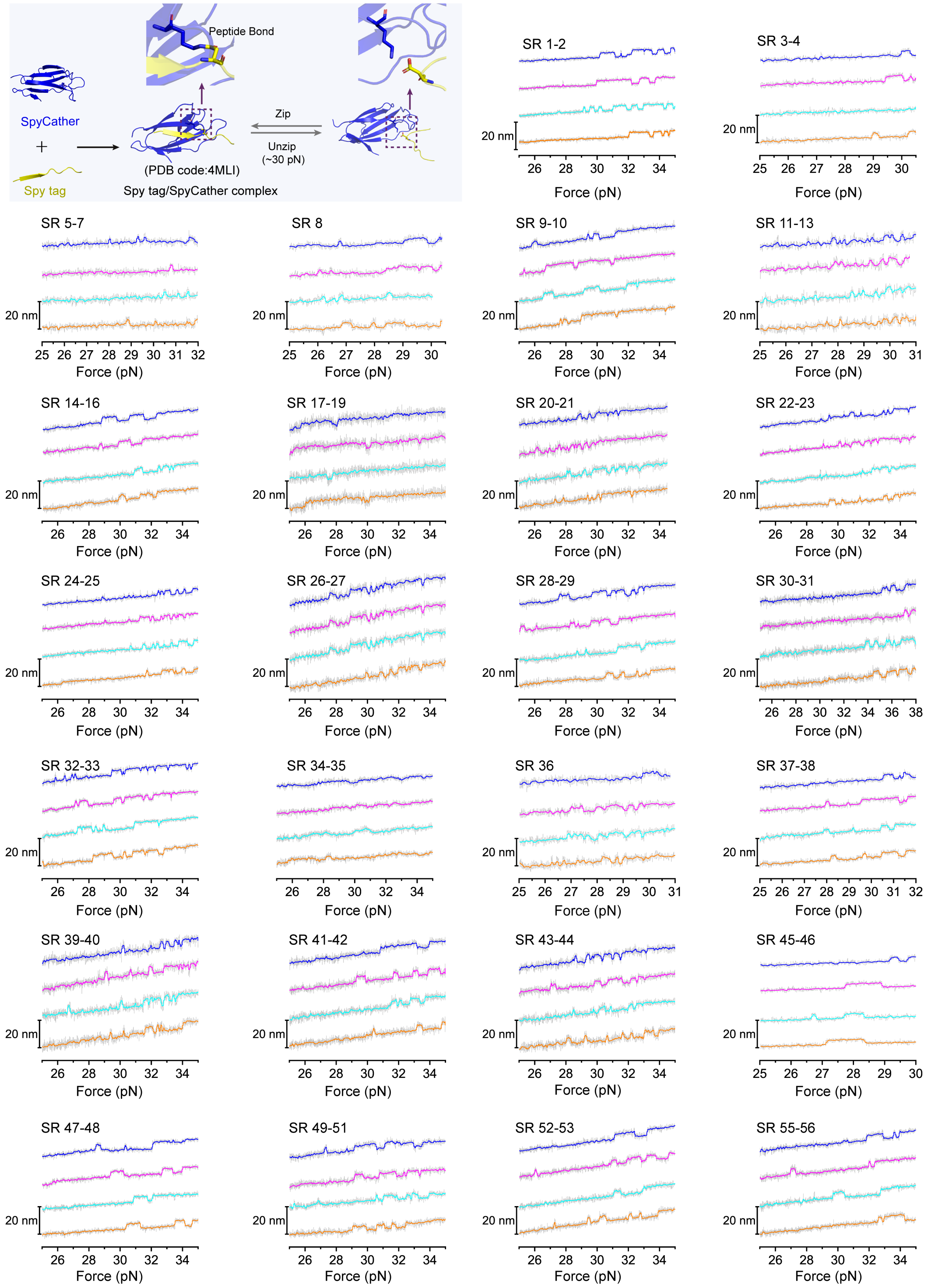


**Supplementary Figure** 1. **The fingerprint signals for all the nesprin-2 SR domain constructs.** Top left panel: The illustration (plotted based on PDB:4MLI) of the unzipping and re-zipping of the SpyTag–SpyCatcher complex, resulted from stretching at the N-terminus of the SpyTag and the N-terminus of the SpyCatcher. The rest of panels: Representative force–bead height curves of all the nesprin-2 SR domain constructs at forces around 30 pN. ~ 4 nm unzipping and re-zipping steps were always observed. The colored lines are 10-point FFT-smooth of the raw data in gray.

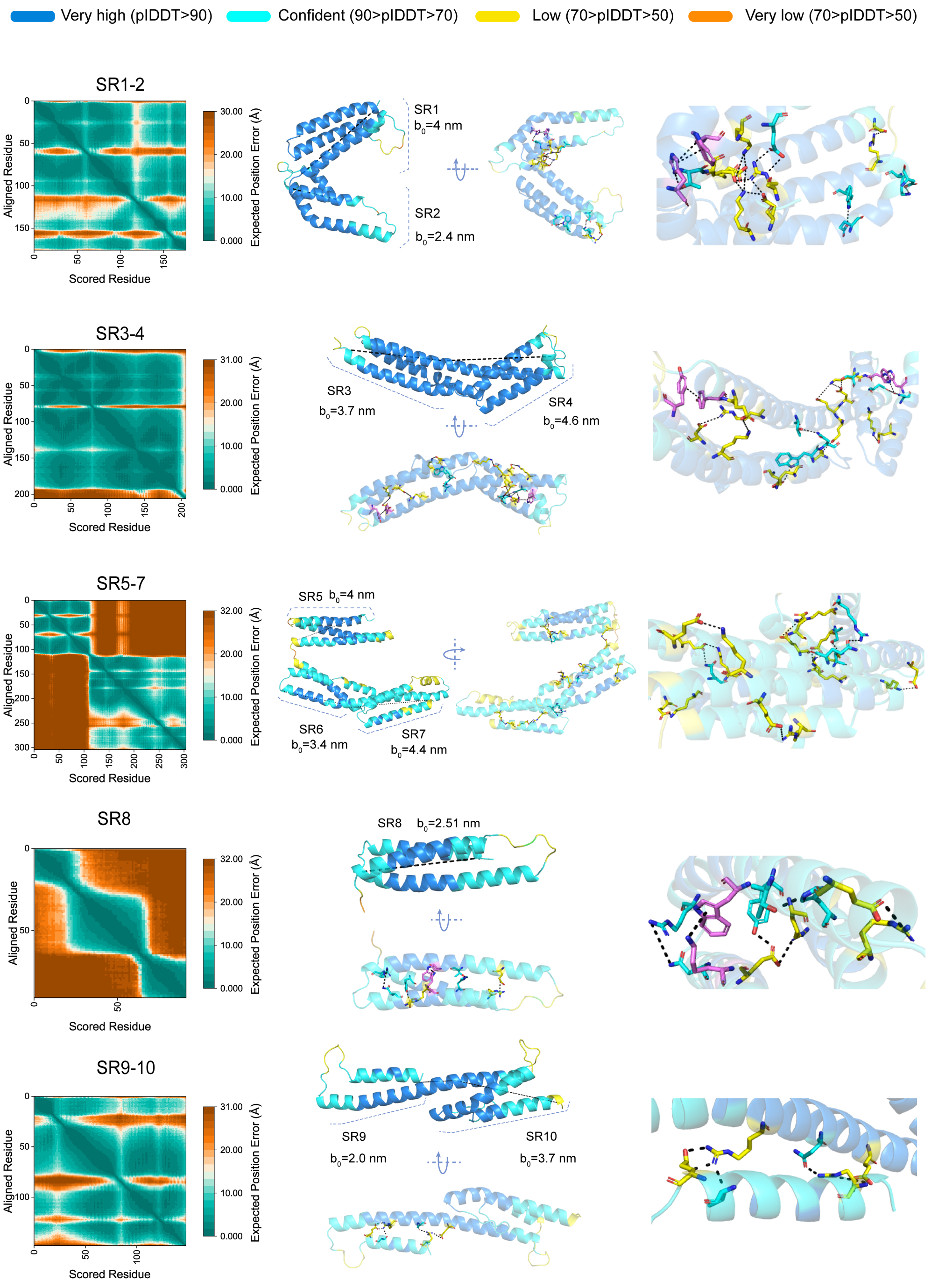


**Supplementary Figure 2. AlphaFold2 Predicted structures of giant nesprin-2 SR1-SR10 domains.** Predicted structures of giant nesprin-2 SR1-SR10 domains generated by AlphaFold2 (AF2). Left panel: the predicted aligned error (PAE) associated with AF2's model 0 predictions. Middle panel: the predicted structure colored with predicted local distance difference test (plDDT) scores, from two different viewing angles. The distance between the N-terminus and the C-terminus of the rigid folded structure (i.e., $b_0$) is measured and labeled. Right panel: the inter-helix interactions analyzed based on the predicted structures, where the $\pi - \pi$ bond, salt-bridge and hydrogen bond are denoted in magenta, yellow and cyan, respectively.

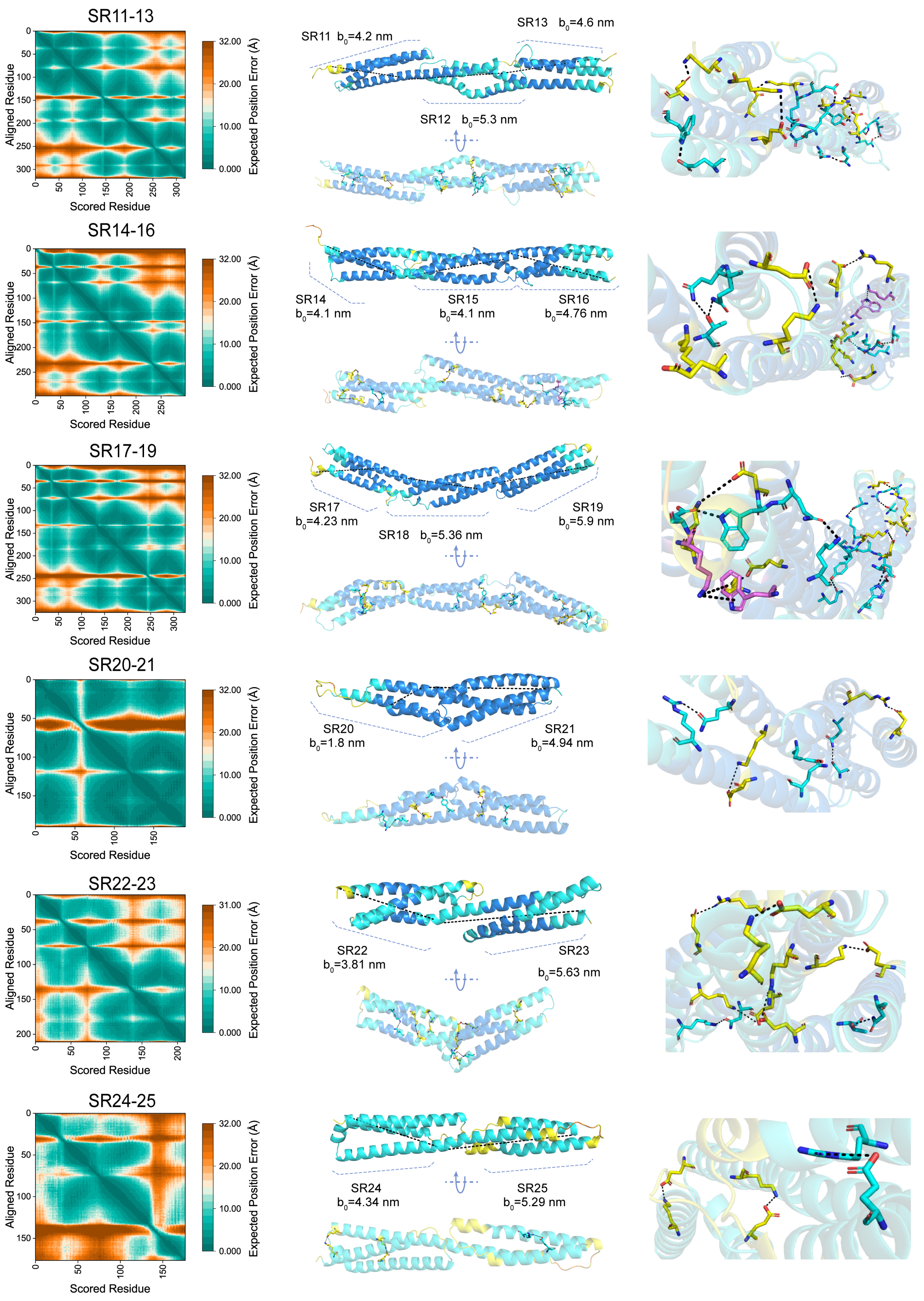


**Supplementary Figure** 3. **AlphaFold2 Predicted structures of giant nesprin-2 SR11-SR25 domains.** Predicted structures of giant nesprin-2 SR1-SR10 domains generated by AlphaFold2 (AF2). Left panel: the predicted aligned error (PAE) associated with AF2's model 0 predictions. Middle panel: the predicted structure colored with predicted local distance difference test (plDDT) scores, from two different viewing angles. The distance between the N-terminus and the C-terminus of the rigid folded structure (i.e., $b_0$) is measured and labeled. Right panel: the inter-helix interactions analyzed based on the predicted structures, where the $\pi - \pi$ bond, salt-bridge and hydrogen bond are denoted in magenta, yellow and cyan, respectively.

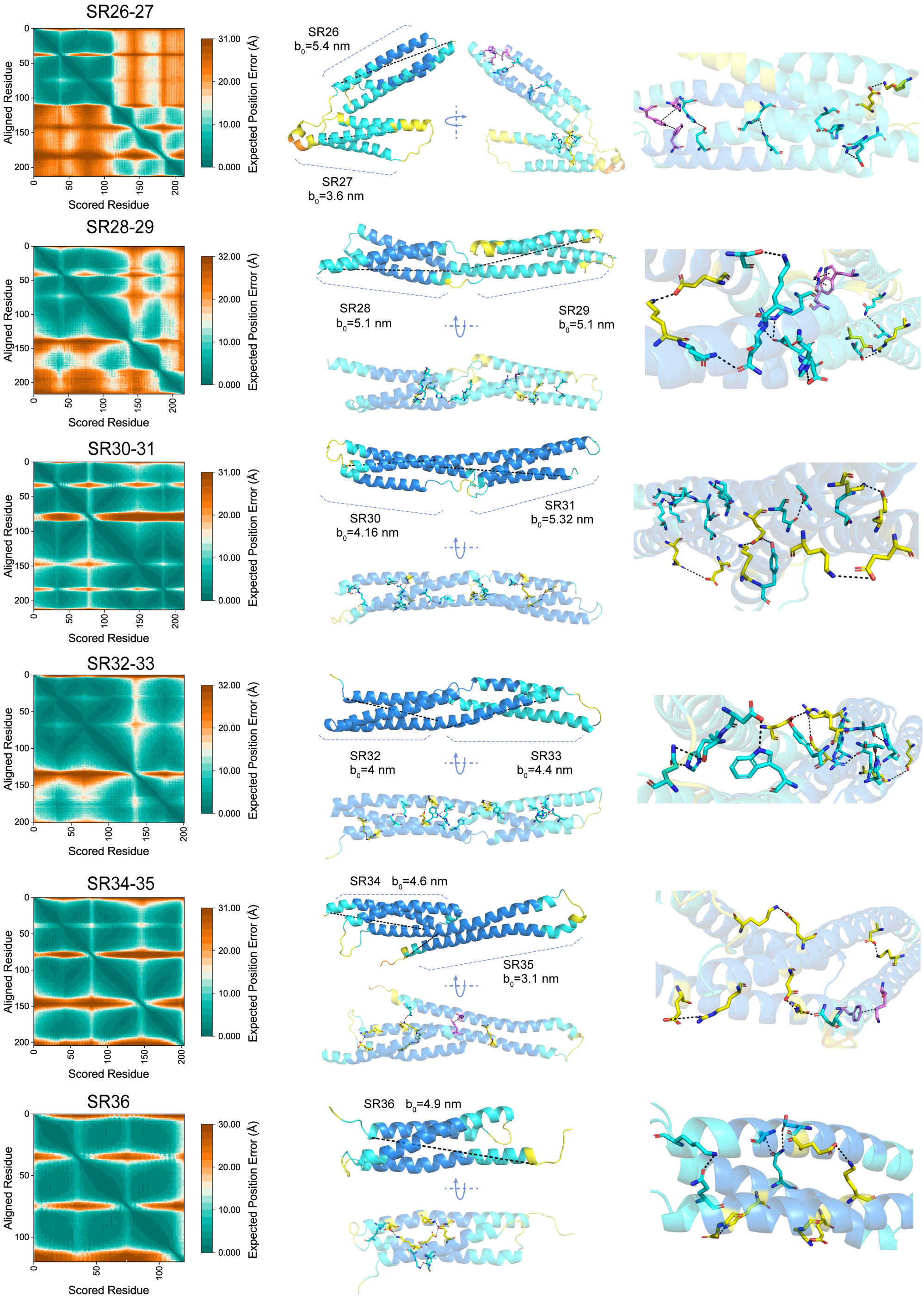


**Supplementary Figure** 4. **AlphaFold2 Predicted structures of giant nesprin-2 SR26-SR36 domains.** Predicted structures of giant nesprin-2 SR1-SR10 domains generated by AlphaFold2 (AF2). Left panel: the predicted aligned error (PAE) associated with AF2's model 0 predictions. Middle panel: the predicted structure colored with predicted local distance difference test (plDDT) scores, from two different viewing angles. The distance between the N-terminus and the C-terminus of the rigid folded structure (i.e., $b_0$) is measured and labeled. Right panel: the inter-helix interactions analyzed based on the predicted structures, where the $\pi - \pi$ bond, salt-bridge and hydrogen bond are denoted in magenta, yellow and cyan, respectively.

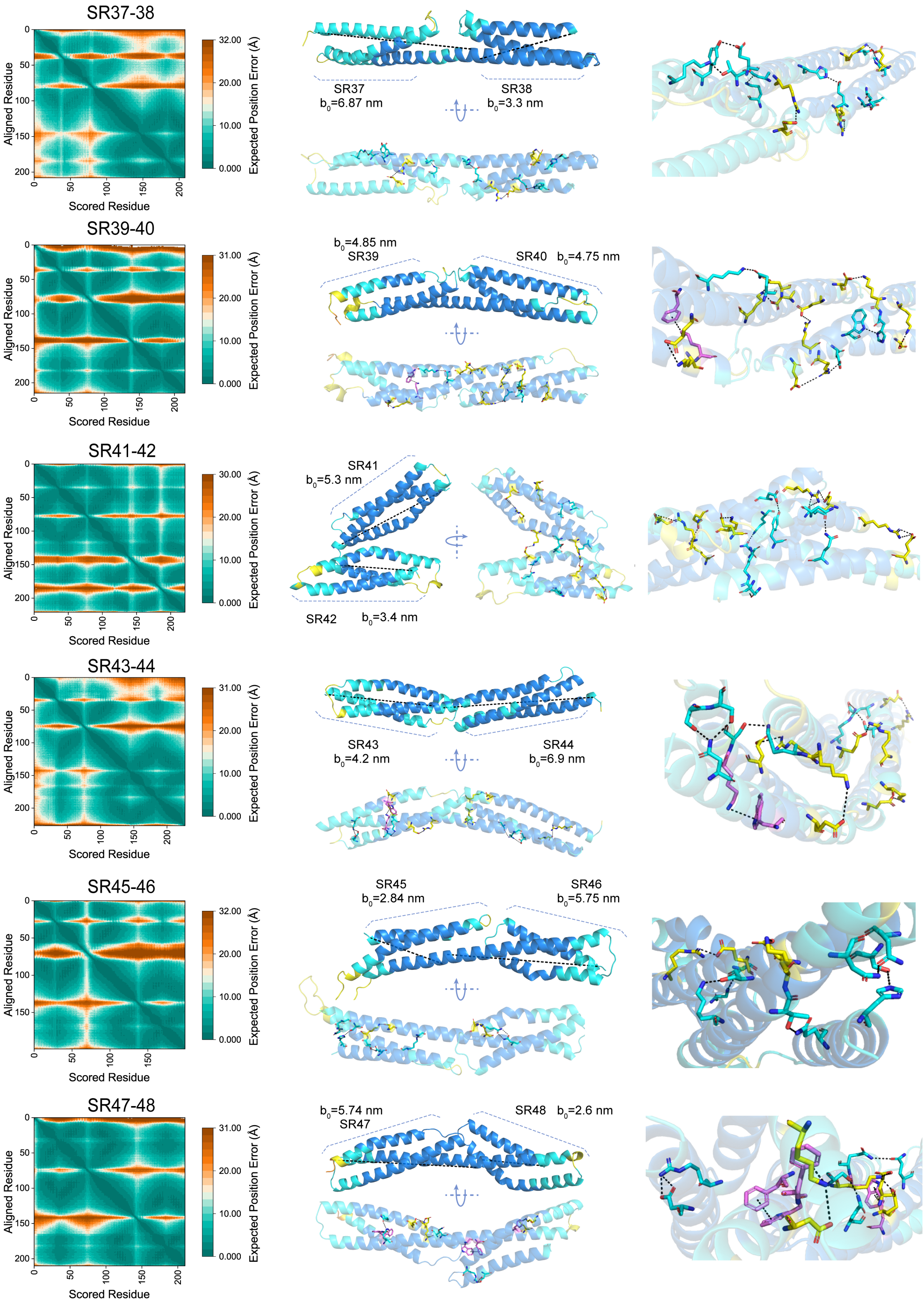


**Supplementary Figure** 5. **AlphaFold2 Predicted structures of giant nesprin-2 SR37-SR48 domains.** Predicted structures of giant nesprin-2 SR1-SR10 domains generated by AlphaFold2 (AF2). Left panel: the predicted aligned error (PAE) associated with AF2's model 0 predictions. Middle panel: the predicted structure colored with predicted local distance difference test (plDDT) scores, from two different viewing angles. The distance between the N-terminus and the C-terminus of the rigid folded structure (i.e., $b_0$) is measured and labeled. Right panel: the inter-helix interactions analyzed based on the predicted structures, where the $\pi - \pi$ bond, salt-bridge and hydrogen bond are denoted in magenta, yellow and cyan, respectively.

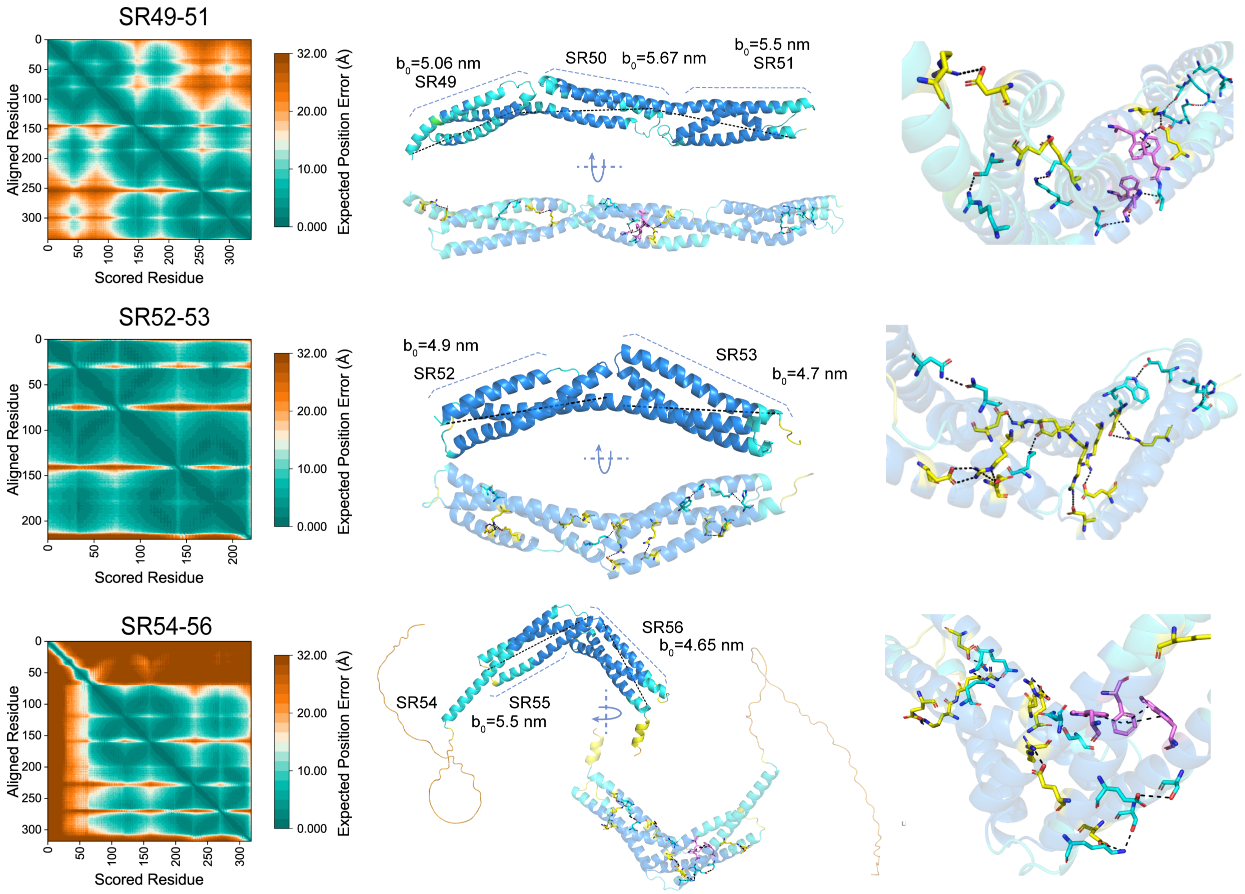


**Supplementary Figure** 6. **AlphaFold2 Predicted structures of giant nesprin-2 SR49-SR56 domains.** Predicted structures of giant nesprin-2 SR1-SR10 domains generated by AlphaFold2 (AF2). Left panel: the predicted aligned error (PAE) associated with AF2's model 0 predictions. Middle panel: the predicted structure colored with predicted local distance difference test (plDDT) scores, from two different viewing angles. The distance between the N-terminus and the C-terminus of the rigid folded structure (i.e., $b_0$) is measured and labeled. Right panel: the inter-helix interactions analyzed based on the predicted structures, where the $\pi - \pi$ bond, salt-bridge and hydrogen bond are denoted in magenta, yellow and cyan, respectively.

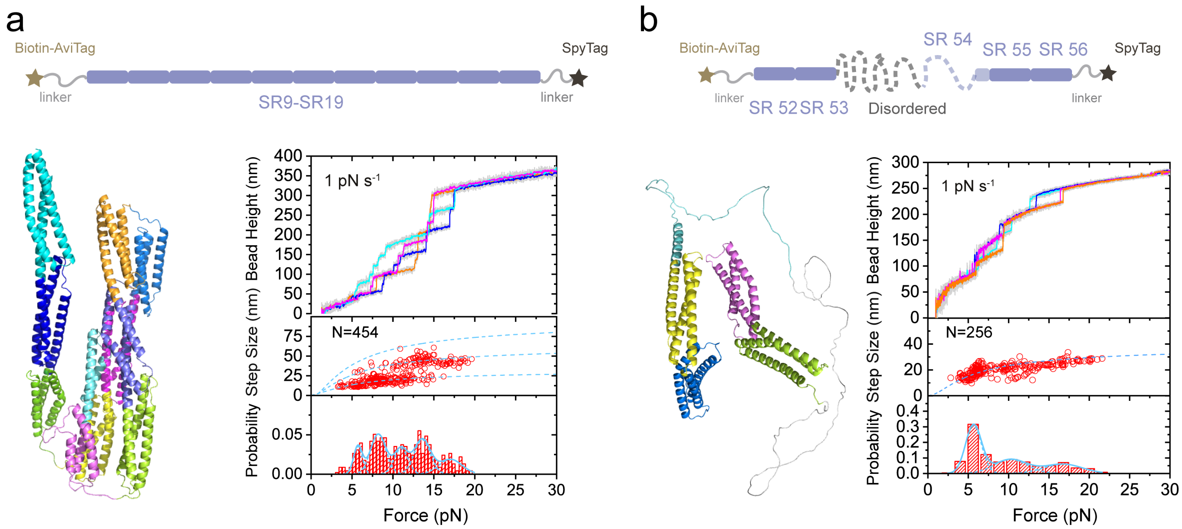


**Supplementary Figure** 7. **The force-responses of the nesprin-2 SR9-SR19 and SR52-SR56.** (**a-b**). Top panel: illustration of the single-molecule construct design for SR9-SR19 (a) and SR52-SR56 (b), respectively. Left panel: The AlphaFold2 predicted structrues of SR9-SR19 (a) and SR52-SR56 (b), respectively. Right Panel: The representative force–bead height curves during linear force-increase scan (1 pN $s^{-1}$), the resulting force–step size curves and the normalized unfolding force distributions for SR9-SR19 (a) and SR52-SR56 (b), respectively. Raw data (gray) is 10-point FFT smoothed (colored lines). Number ($N$) of data points (represented by hollow circles) are indicated in the panels. The dashed colored lines are theoretical calculation of the force-dependent unfolding/refolding step sizes for corresponding SR domains. The colored solid lines are multiple Gaussian fitting of the distributions.

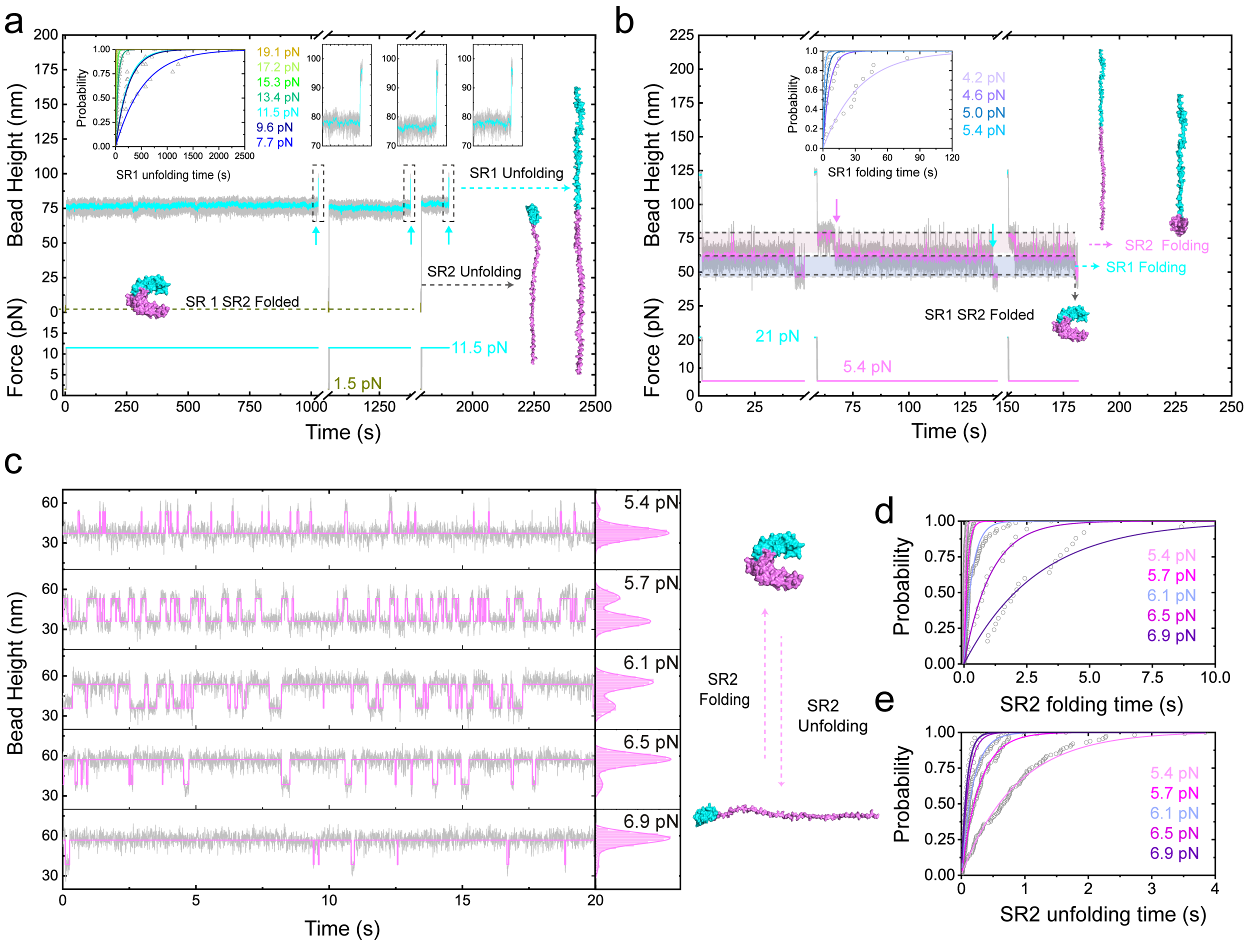


**Supplementary Figure** 8. **Force-dependent lifetimes of the giant nesprin-2 SR1-SR2**. (**a**) Representative time–bead height curves of the SR1-SR2 during force-jump scans from 1 pN to 11.5 pN (cyan). The upward arrows indicate the unfolding events of SR1 at 11.5 pN. The SR2 unfolds during force-increase from 1 pN to 11.5 pN. The inset panel: The resulting unfolding time probability of the SR1 at forces of 7-20 pN. The lines are fitting curves of the probability distribution. (**b**) Representative time–bead height curves of the SR1-SR2 during force-jump scans from 21 pN to 5.4 pN (magenta). The downward arrows indicate the refolding events of SR2 (magenta arrow) and SR1 (blue arrow) at 5.4 pN. The inset panel: The resulting refolding time probability of the SR1 at forces of 4-5.4 pN. The lines are fitting curves of the probability distribution. (**c**) Representative time–bead height curves of the SR1-SR2 during constant-force measurement at forces of 5-7 pN. Unfolding/refolding dyanmics of SR2 at these forces are analyzed. The raw data (gray) is 10-point FFT smoothed (colored lines). (**d-e**) The resulting folding (d) and unfolding (e) time probability of the SR2 at forces of 4-7 pN. The lines are corresponding fitting curves of the probability distribution.

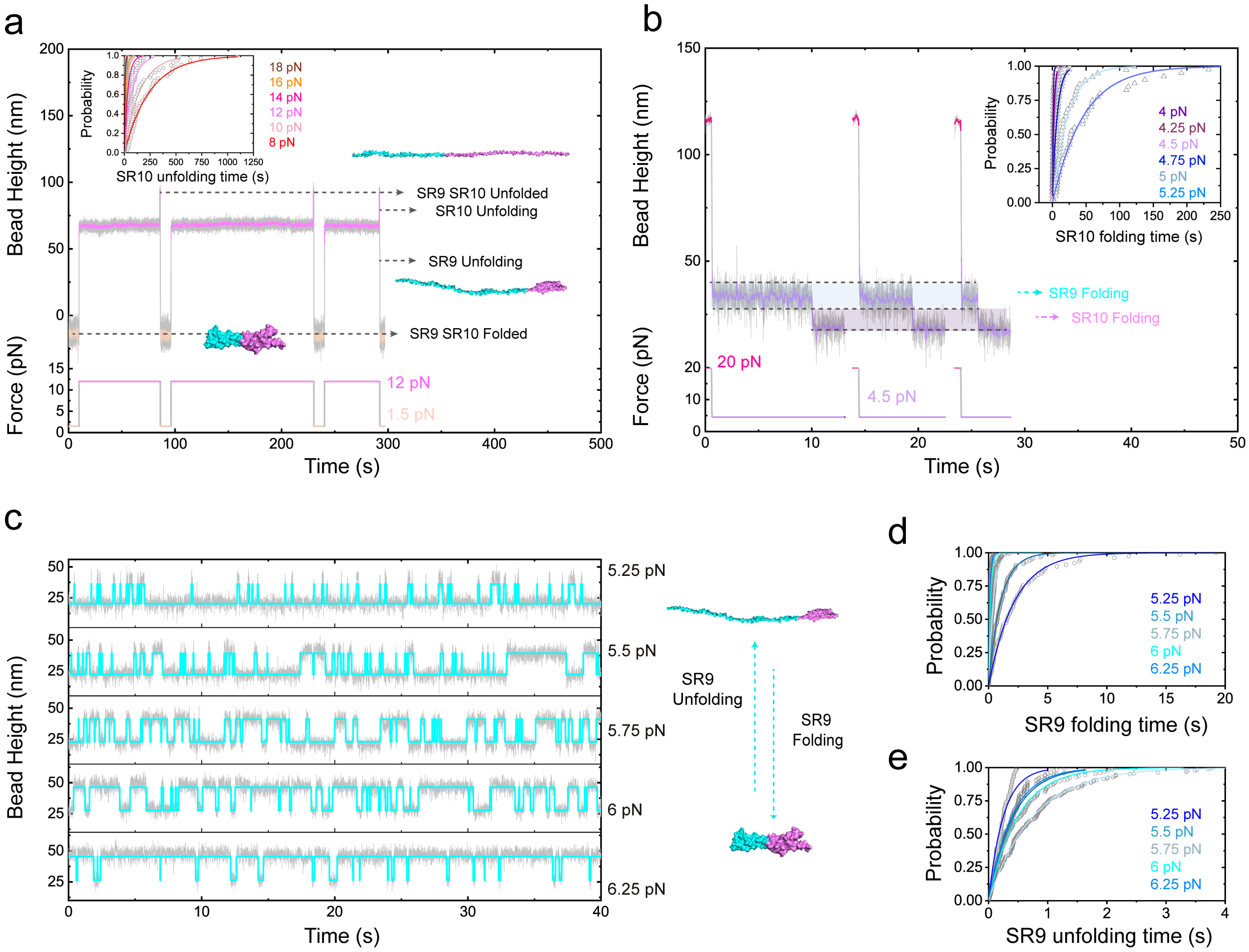


**Supplementary Figure 9. Force-dependent lifetimes of the giant nesprin-2 SR9-SR10.** (**a**) Representative time–bead height curves of the SR9-SR10 during force-jump scans from 1.5 pN to 12 pN (magenta). The SR9 unfolds during force-increase. The inset panel: The resulting unfolding time probability of the SR10 at forces of 8-18 pN. The lines are fitting curves of the probability distribution. (**b**) Representative time–bead height curves of the SR9-SR10 during force-jump scans from 20 pN to 4.5 pN (purple). The inset panel: The resulting refolding time probability of the SR1 at forces of 4-5.4 pN. The lines are fitting curves of the probability distribution. (**c**) Representative time–bead height curves of the SR9-SR10 during constant-force measurement at forces of 5-6 pN. Unfolding/refolding dyanmics of SR9 at these forces are analyzed. The raw data (gray) is 10-point FFT smoothed (colored lines). (**d-e**) The resulting folding (d) and unfolding (e) time probability of the SR9 at forces of 5-6 pN. The lines are corresponding fitting curves of the probability distribution.

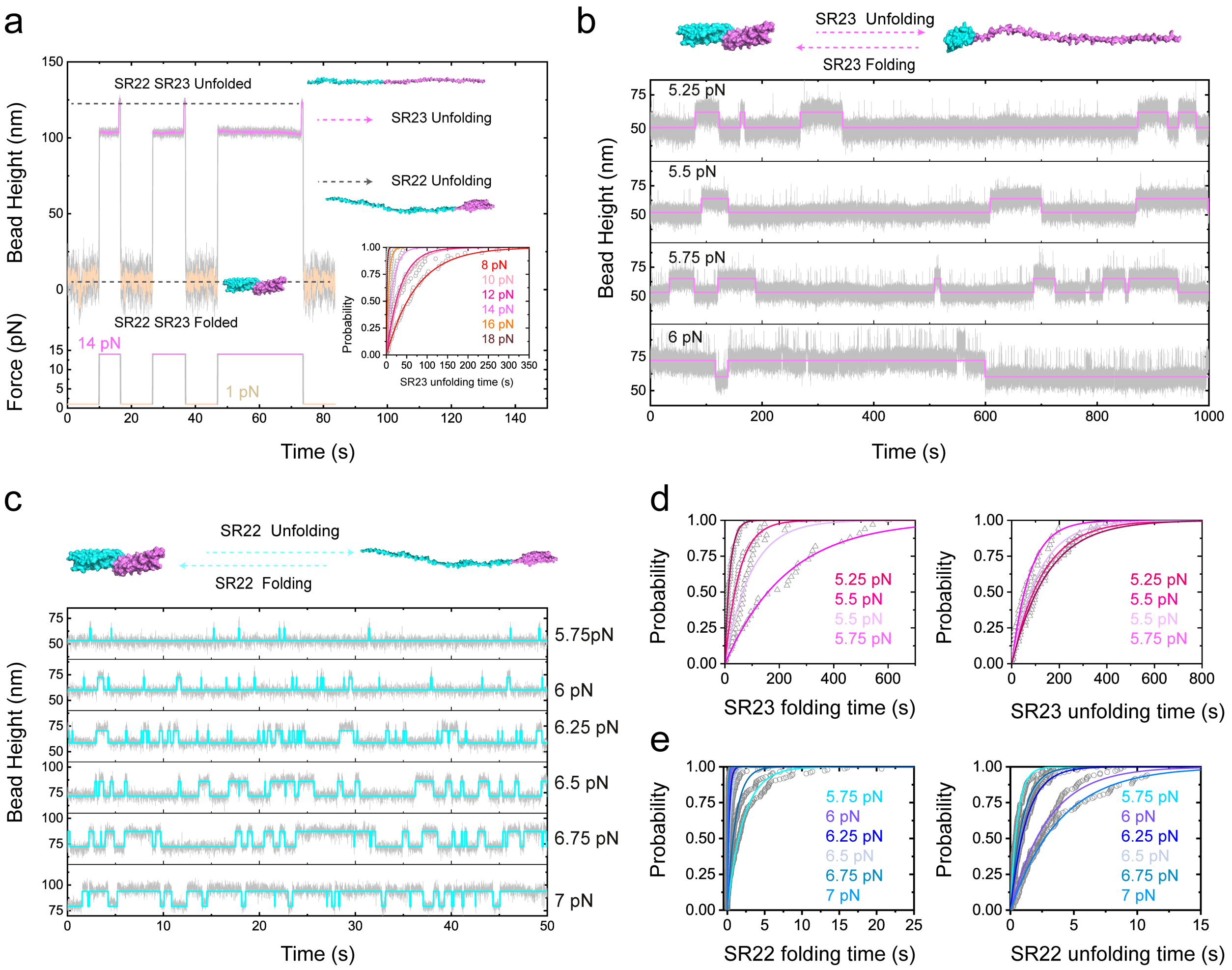


**Supplementary Figure** 10. **Force-dependent lifetimes of the giant nesprin-2 SR22-SR23**. (**a**) Representative time–bead height curves of the SR22-SR23 during force-jump scans from 1 pN to 14 pN (magenta). The inset panel: The resulting unfolding time probability of the SR1 at forces of 8-18 pN. The lines are fitting curves of the probability distribution. (**b-c**) Representative time–bead height curves of the SR22-SR23 during constant-force measurement at forces of 5-7 pN. Unfolding/refolding dynamics of SR23 (b) and SR22 (c) at these forces are analyzed. The raw data (gray) is 10-point FFT smoothed (colored lines). (**d-e**) The resulting folding (d) and unfolding (e) time probability of SR22 and SR23 at forces of 4-7 pN. The lines are corresponding fitting curves of the probability distribution.

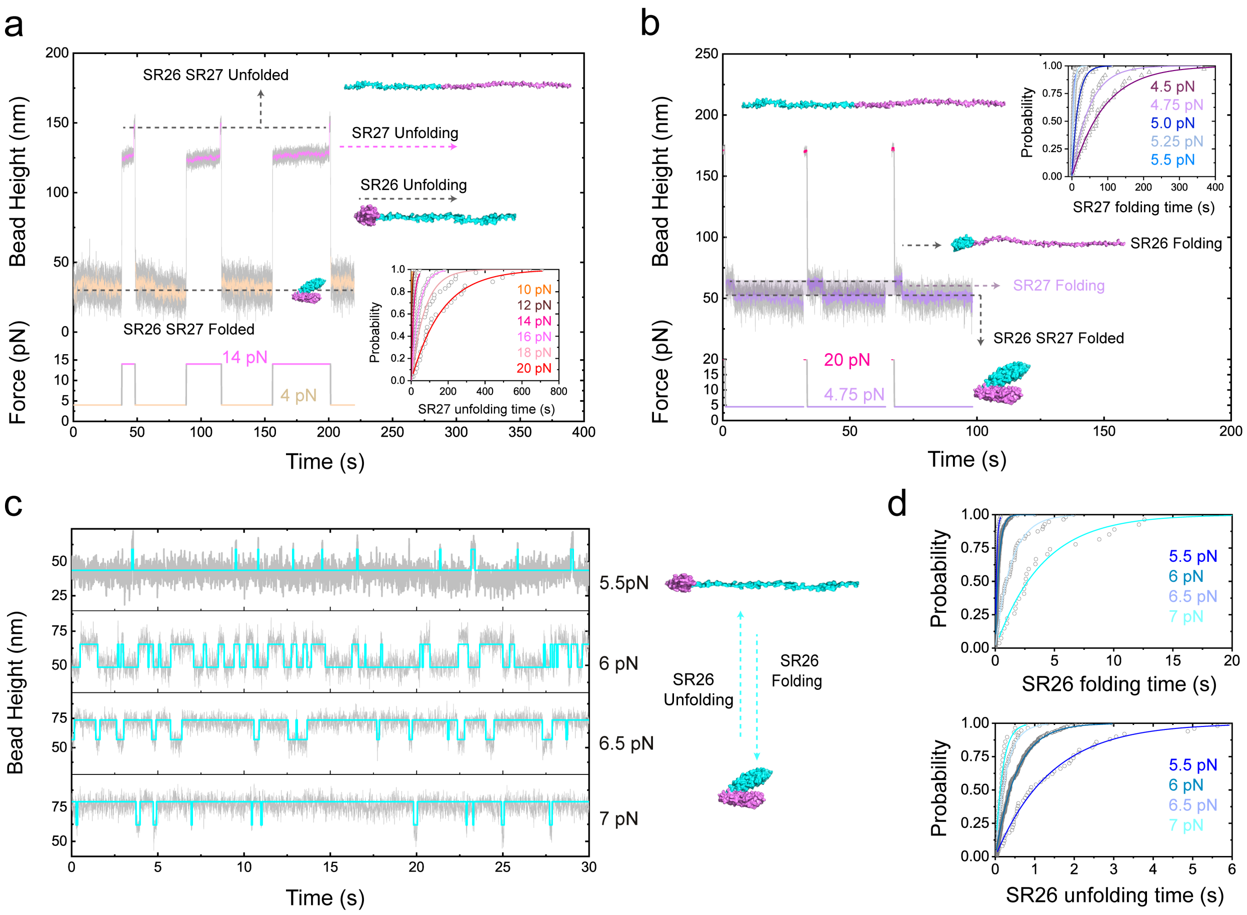


**Supplementary Figure 11. Force-dependent lifetimes of the giant nesprin-2 SR26-SR27.** (**a**) Representative time–bead height curves of the SR26-SR27 during force-jump scans from 4 pN to 14 pN (magenta). The inset panel: The resulting unfolding time probability of the SR27 at forces of 10-20 pN. The lines are fitting curves of the probability distribution. (**b**) Representative time–bead height curves of the SR26-SR27 during force-jump scans from 20 pN to 4.75 pN (purple). The inset panel: The resulting refolding time probability of the SR27 at forces of 4.5-5.5 pN. The lines are fitting curves of the probability distribution. (**c**) Representative time–bead height curves of the SR26-SR27 during constant-force measurement at forces of 5-7 pN. Unfolding/refolding dynamics of SR26 at these forces are analyzed. The raw data (gray) is 10-point FFT smoothed (colored lines). (**d**) The resulting folding and unfolding time probability of SR26 at forces of 5-7 pN. The lines are corresponding fitting curves of the probability distribution.

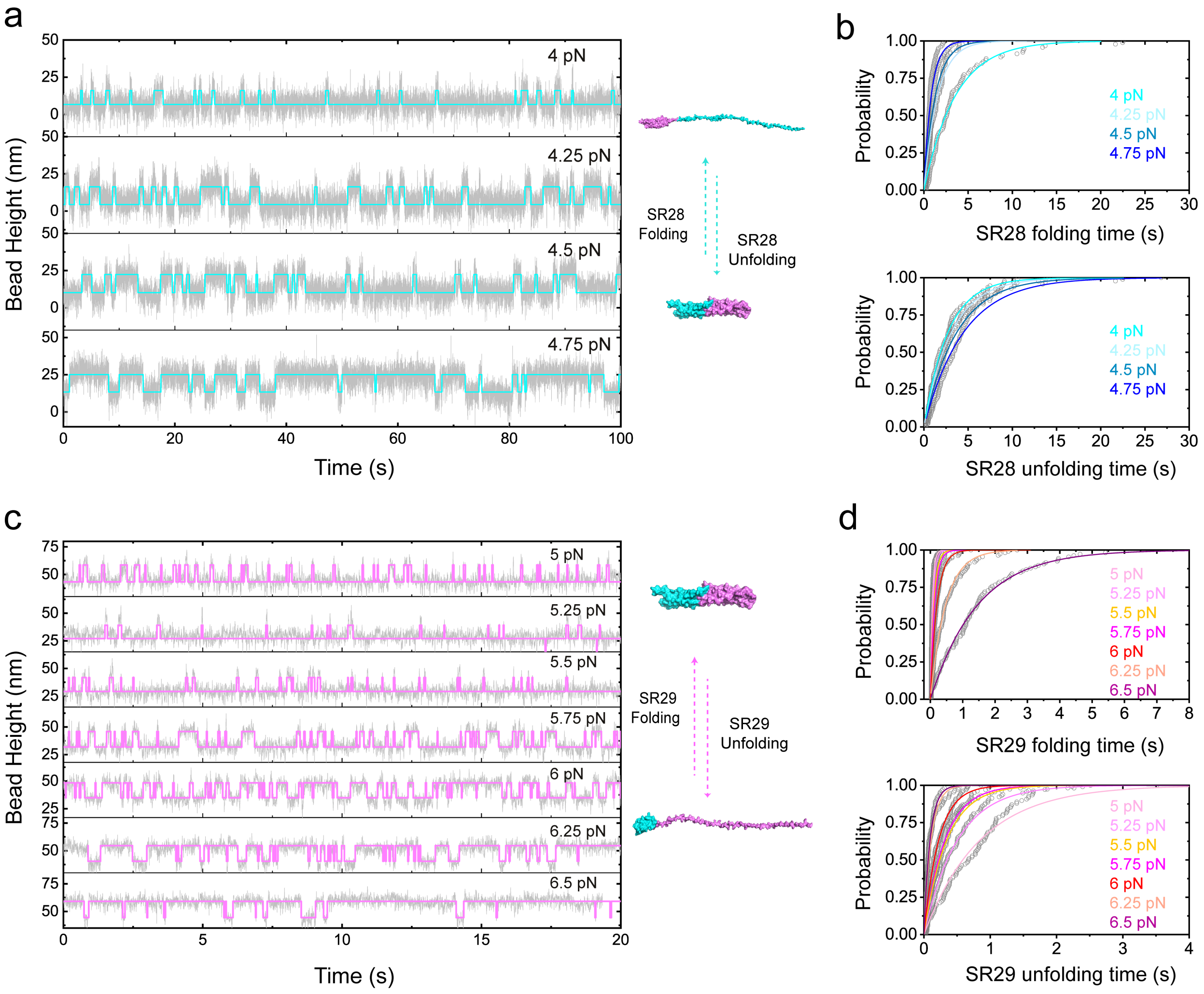


**Supplementary Figure** 12. **Force-dependent lifetimes of the giant nesprin-2 SR28-SR29**. (**a-d**) Representative time–bead height curves of the SR28-SR29 during constant-force measurement at forces of 4-7 pN. Unfolding/refolding dynamics of SR28 (a) and SR29 (c) at these forces are analyzed. The raw data (gray) is 10-point FFT smoothed (colored lines). (**b**) The resulting folding and unfolding time probability of SR28.(**d**) The folding and unfolding time probability of SR29, at forces of 5-7 pN are fitted (colored lines).

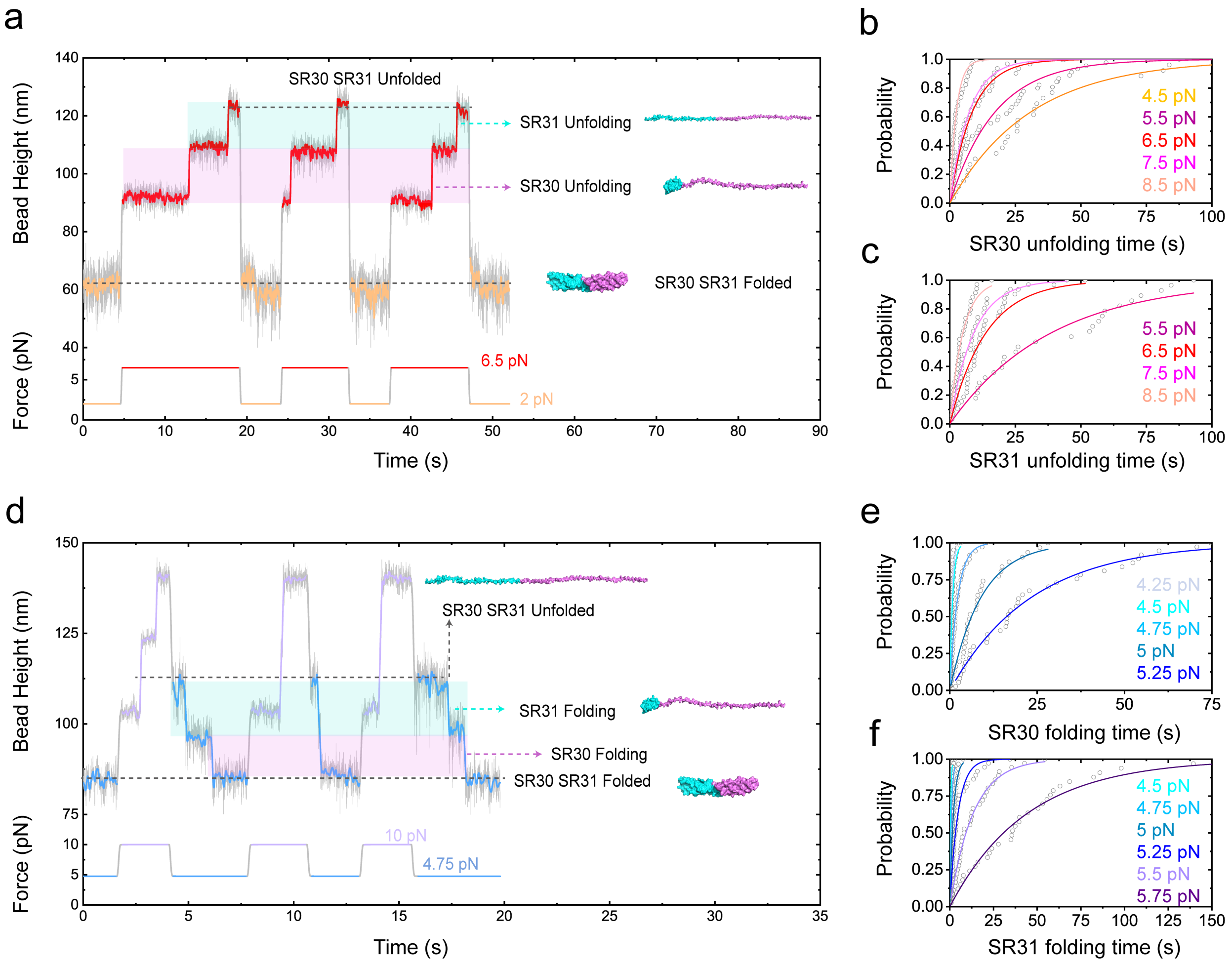


**Supplementary Figure** 13. **Force-dependent lifetimes of the giant nesprin-2 SR30-SR31**. (**a**) Representative time–bead height curves of the SR30-SR31 during force-jump scans from 2 pN to 6.5 pN (red). (**b**) The resulting unfolding time probability of the SR30 (b) and SR31 (c) at forces of 4-8 pN. The lines are fitting curves of the probability distribution. (**c**) Representative time–bead height curves of the SR30-SR31 during force-jump scans from 10 pN to 4.75 pN (blue). (**e-f**) The resulting refolding time probability of the SR30 (e) and SR31 (f) at forces of 4-5 pN. The lines are fitting curves of the probability distribution.

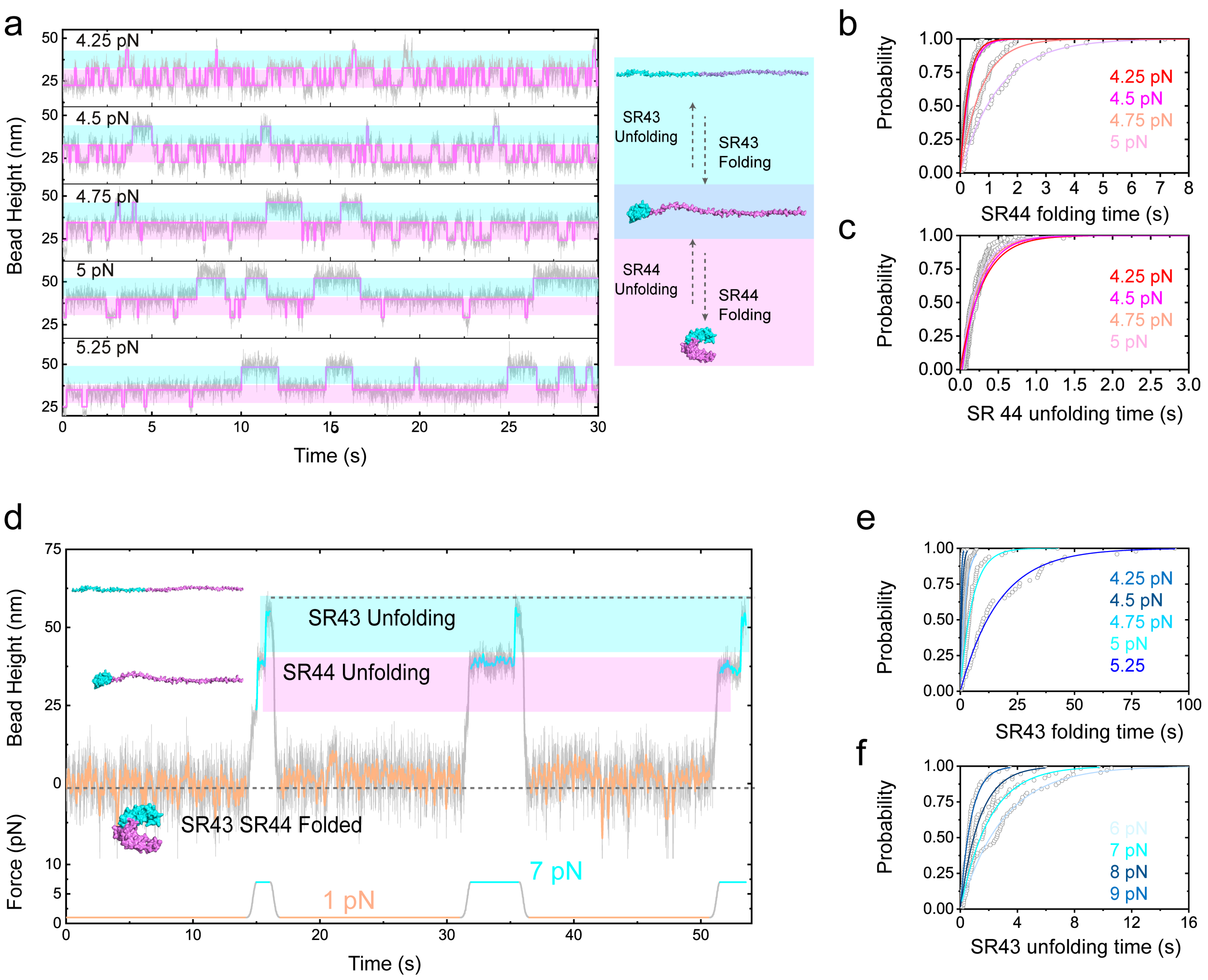


**Supplementary Figure** 14. **Force-dependent lifetimes of the giant nesprin-2 SR43-SR44**. (**a**) Representative time–bead height curves of the SR43-SR44 during constant-force measurement at forces of 4-5 pN. Unfolding/refolding dynamics of SR43 (light blue) and SR44 (light magenta) at these forces are analyzed. The raw data (gray) is 10-point FFT smoothed (colored lines). (**b-c**) The resulting folding (b) and unfolding (c) time probability of SR44 are fitted (colored lines). (**d**) Representative time–bead height curves of the SR43-SR44 during constant-force measurement at forces of 5-6 pN. Unfolding/refolding dynamics of SR43 (cyan) at these forces are analyzed. The raw data (gray) is 10-point FFT smoothed (colored lines). (**e**) Representative time–bead height curves of the SR43-SR44 during force-jump scans from 1 pN to 7 pN (cyan). (**f**) The resulting folding (top) and unfolding (bottom) time probability of SR43 are fitted (colored lines).

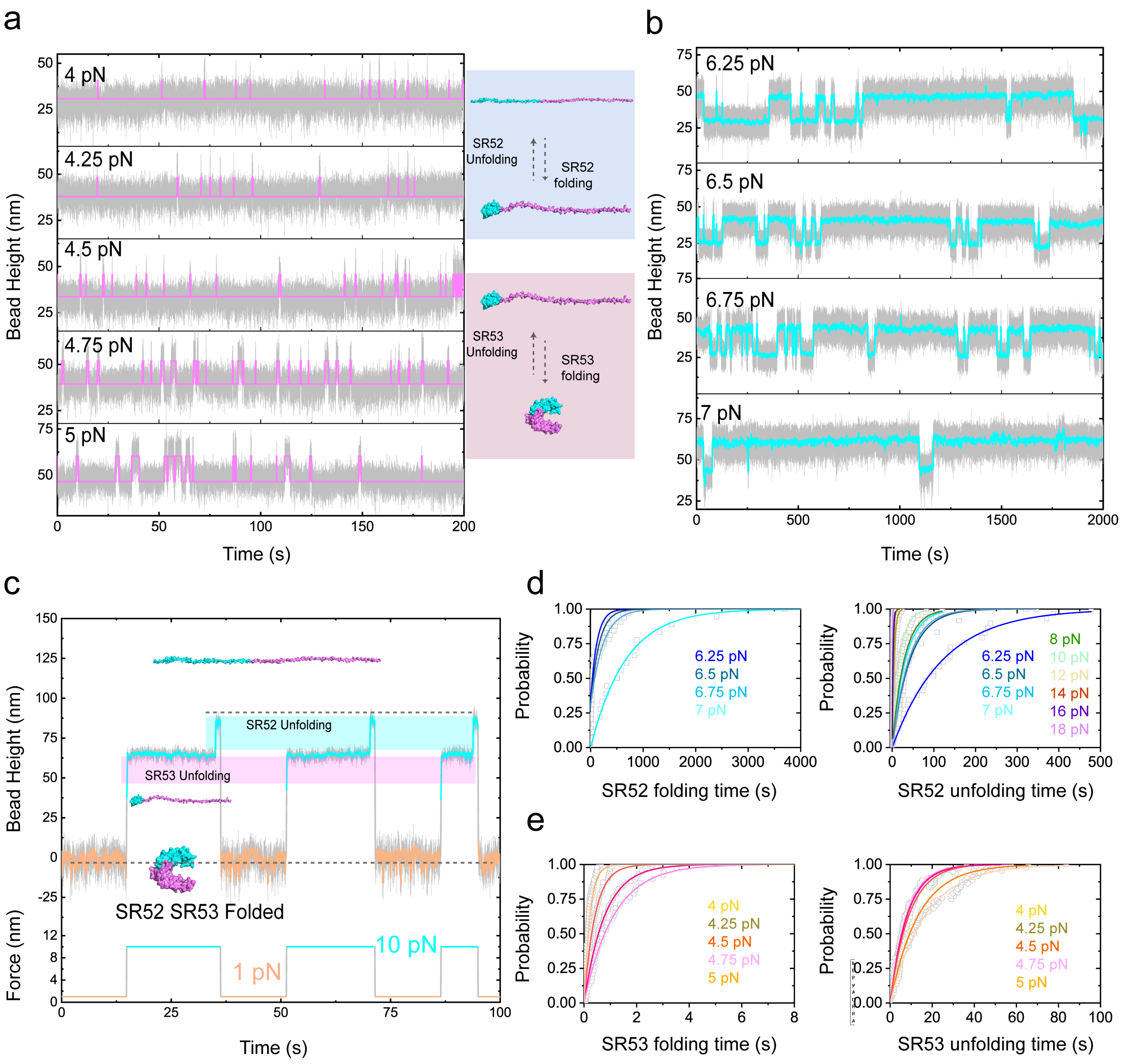


**Supplementary Figure** 15. **Force-dependent lifetimes of the giant nesprin-2 SR52-SR53**. (**a**) Representative time–bead height curves of the SR52-SR53 during constant-force measurement at forces of 4-5 pN. The raw data (gray) is 10-point FFT smoothed (colored lines). (**b-c**) The resulting folding (b) and unfolding (c) time probability of SR53 are fitted (colored lines). (**d**) Representative time–bead height curves of the SR52-SR53 during constant-force measurement at forces of 4-6 pN. Unfolding/refolding dynamics of SR52 (cyan) at these forces are analyzed. The raw data (gray) is 200-point FFT smoothed (cyan lines). (**e**) Representative time–bead height curves of the SR52-SR53 during force-jump scans from 8 pN to 18 pN (cyan). (**f**) The resulting folding (top) and unfolding (bottom) time probability of SR52 are fitted (colored lines).

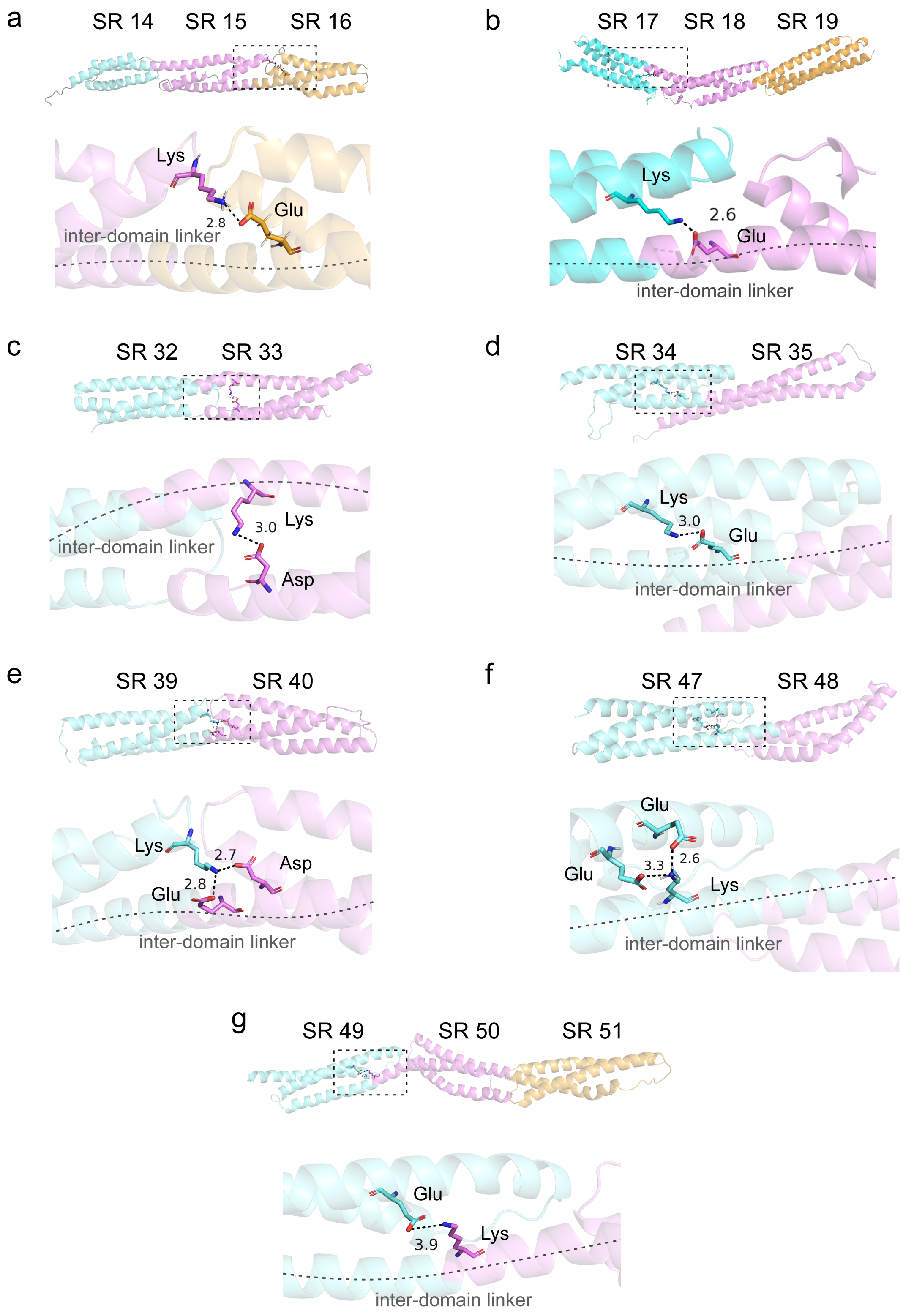


**Supplementary Figure** 16. **The inter-domain salt-bridges within the SR domains.** (**a-g**). The inter-domain salt-bridges within the SR domains of the giant nesprins analyzed based on the AlphaFold2 predicted structures. The concurrent unfolding neighboring SR domains are analyzed.

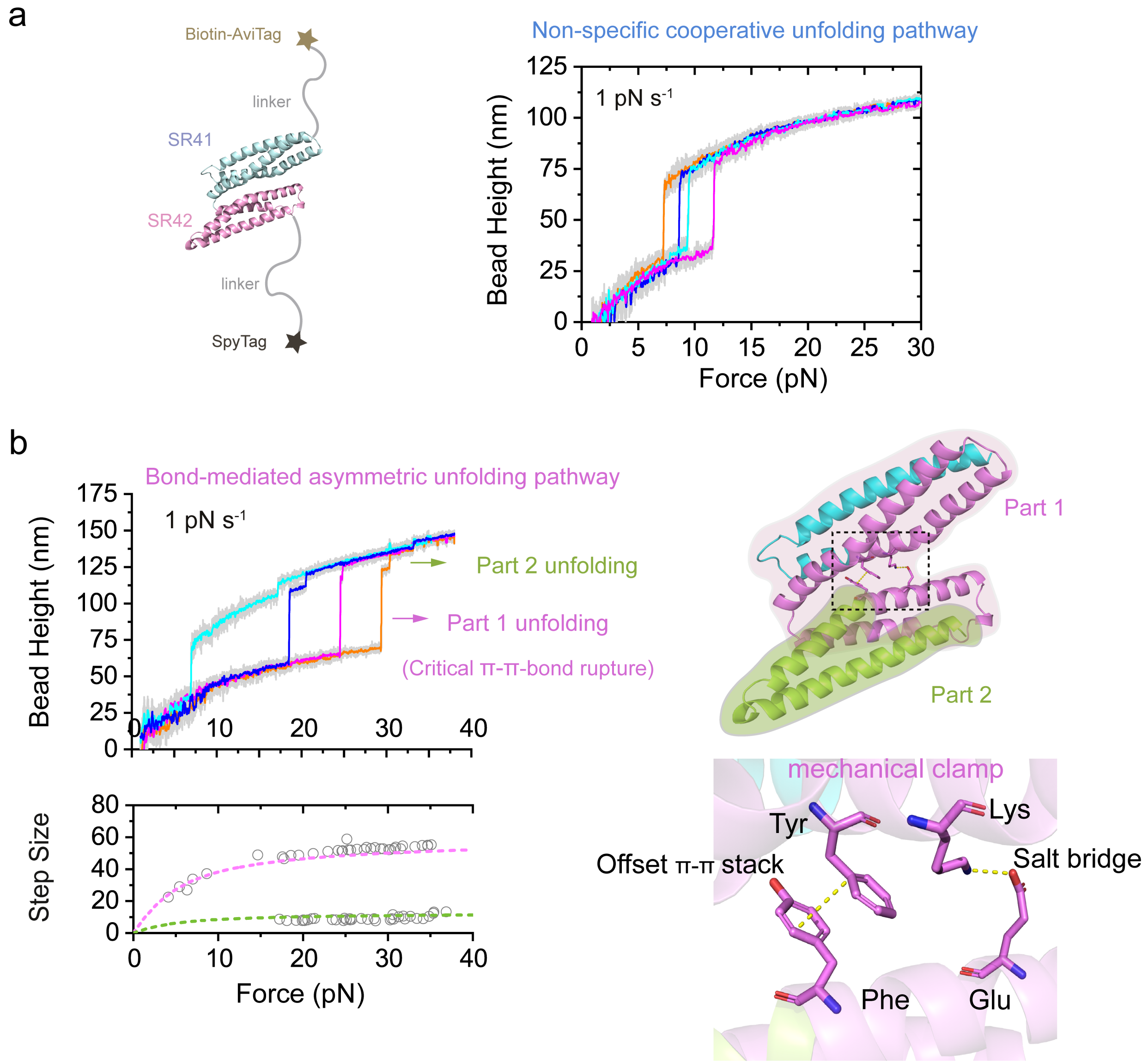


**Supplementary Figure** 17. $\pi - \pi$ **bond-mediated unfolding pathway**. (**a**) Left panel: illustrations of single-molecule manipulation of SR41-42. Right panel: Representative force–bead height curves of SR41-42 during linear force-increase scans 1 pN s$^{-1}$ reveal cooperative unfolding pathways mediated by non-specific strong interactions. (**b**) Left panel: Representative force–bead height curves of the SR 41-42 during linear force-increase 1 pN s$^{-1}$ scans showing domain-specific unfolding, where the unfolding step sizes quantitatively agree with theoretical predictions for the sequential rupture of Part 1 (magenta dash) and Part 2 (green dash) domains following $\pi{-}\pi$ stacking disruption. Mechanical disruption pathway: external force preferentially breaks the parallel $\pi - \pi$ stacking between residues Tyr and Phe, leading to subsequent salt bridge dissociation and unilateral domain unfolding (Part 1). Right panel: In the native conformation, residues Tyr and Phe form an offset $\pi - \pi$ stacking interaction, which cooperates with a salt bridge network (e.g., Lys and Glu) to jointly stabilize the domain interface. Bottom panel: Detailed view of the mechanical clamp region, highlighting the offset $\pi - \pi$ stack and salt bridge interactions that contribute to domain stability.

| Domain | $k_u^0$ ($s^{-1}$) | $x_u$ (nm) | $k_f^0$ ($s^{-1}$) | $b^\star$(nm) |
|---|---|---|---|---|
| SR1 | $2.96\times10^{-5}$ | 2.01 | $9.57\times10^{2}$ | $5.01\times10^{-3}$ |
| SR2 | $5.36\times10^{-5}$ | 8.23 | $5.31\times10^{2}$ | 7.06 |
| SR3 | $6.09\times10^{-4}$ | 7.11 | $1.78\times10^{2}$ | 5.03 |
| SR4 | $6.09\times10^{-4}$ | 7.11 | $1.78\times10^{2}$ | 5.03 |
| SR5 | $3.03\times10^{-4}$ | 2.26 | $1.15\times10^{3}$ | 8.18 |
| SR6 | $4.73\times10^{-3}$ | 1.36 | $1.15\times10^{3}$ | 8.18 |
| SR7 | $7.81\times10^{-4}$ | 5.37 | $3.52\times10^{2}$ | 4.08 |
| SR8 | $1.60\times10^{-2}$ | 1.72 | 0.38 | 8.05 |
| SR9 | $2.96\times10^{-4}$ | 5.27 | 73.62 | 9.38 |
| SR10 | $1.07\times10^{-4}$ | 1.84 | 38.24 | 4.79 |
| SR11 | $4.46\times10^{-4}$ | 2.53 | 79.72 | $5.12\times10^{-5}$ |
| SR12 | $2.44\times10^{-3}$ | 2.80 | 0.16 | 10.50 |
| SR13 | $3.48\times10^{-4}$ | 4.39 | $3.51\times10^{2}$ | 5.09 |
| SR14 | $1.34\times10^{-3}$ | 2.50 | $2.28\times10^{2}$ | 7.19 |
| SR15 | $2.07\times10^{-4}$ | 2.48 | $2.28\times10^{2}$ | 7.19 |
| SR16 | $2.07\times10^{-4}$ | 2.48 | 3.30 | 10.45 |
| SR17 | $1.94\times10^{-6}$ | 2.93 | $3.25\times10^{4}$ | 3.56 |
| SR18 | $1.94\times10^{-6}$ | 2.93 | $9.75\times10^{2}$ | 5.39 |
| SR19 | $9.48\times10^{-3}$ | 1.31 | $6.85\times10^{3}$ | 6.07 |
| SR20 | - | - | - | - |
| SR21 | $4.84\times10^{-3}$ | 3.76 | $1.17\times10^{2}$ | 8.16 |
| SR22 | $2.28\times10^{-5}$ | 6.64 | $2.53\times10^{5}$ | 4.95 |
| SR23 | $1.13\times10^{-3}$ | 1.29 | $2.09\times10^{2}$ | 2.98 |
| SR24 | - | - | - | - |
| SR25 | - | - | - | - |
| SR26 | $2.62\times10^{-5}$ | 7.00 | $1.10\times10^{4}$ | 6.21 |
| SR27 | $2.07\times10^{-5}$ | 2.10 | $3.22\times10^{2}$ | 2.20 |
| SR28 | $1.28\times10^{-2}$ | 2.69 | $2.03\times10^{2}$ | 8.28 |
| SR29 | $1.39\times10^{-3}$ | 6.00 | $3.22\times10^{3}$ | 7.00 |
| SR30 | $1.00\times10^{-3}$ | 2.20 | $8.44\times10^{2}$ | 6.72 |
| SR31 | $1.99\times10^{-3}$ | 2.30 | $1.66\times10^{2}$ | 9.48 |
| SR32 | $1.11\times10^{-2}$ | 1.36 | $1.17\times10^{2}$ | 3.86 |
| SR33 | $1.11\times10^{-2}$ | 1.36 | $1.17\times10^{2}$ | 3.86 |
| SR34 | $6.65\times10^{-5}$ | 6.14 | 2.26 | 9.71 |
| SR35 | $6.65\times10^{-5}$ | 6.14 | 2.26 | 9.71 |
| SR36 | $1.96\times10^{-5}$ | 9.54 | 10.31 | 8.77 |
| SR37 | $4.28\times10^{-4}$ | 10.91 | $2.59\times10^{2}$ | 3.41 |
| SR38 | $4.28\times10^{-4}$ | 10.91 | $5.60\times10^{2}$ | 2.41 |
| SR39 | $9.43\times10^{-8}$ | 14.96 | $6.21\times10^{4}$ | $3.25\times10^{-2}$ |

**Supplementary Table** 1. **Kinetic parameters of nesprin SR1-SR39.** The zero-force unfolding rate ($k_u^0$), zero-force refolding rate ($k_f^0$), the unfolding transition distance ($x_u$) and the folding transtion rigid body length ($b^\star$) are listed. The $k_u^0$ and $x_u$ are obtained based on Bell's model fitting to the unfolding force distribution of the domains. The $k_f^0$ and $l^\star$ are obtained by fitting to the refolding force distribution of the domains based on Arrhenius Law. The cooperative unfolding/refolding domains are assigned with same parameters values. The parameters of those domains without clear unfolding/refolding dynamics are not obtained (indicated by '−').

| Domain | $k_u^0$ (s$^{-1}$) | $x_u$ (nm) | $k_f^0$ (s$^{-1}$) | $b^\star$(nm) |
|---|---|---|---|---|
| SR40 | $9.43\times10^{-8}$ | 14.96 | $6.21\times10^{4}$ | $3.25\times10^{-2}$ |
| SR41 | $3.11\times10^{-3}$ | 2.88 | 78.48 | 3.65 |
| SR42 | $3.11\times10^{-3}$ | 2.88 | $3.66\times10^{2}$ | 5.86 |
| SR43 | $8.21\times10^{-3}$ | 2.49 | $1.35\times10^{5}$ | 1.65 |
| SR44 | $2.02\times10^{-4}$ | 8.96 | $1.39\times10^{4}$ | 8.09 |
| SR45 | - | - | - | - |
| SR46 | - | - | - | - |
| SR47 | $3.40\times10^{-6}$ | 8.78 | 22.67 | 3.23 |
| SR48 | $1.61\times10^{-5}$ | 9.98 | 22.67 | 3.23 |
| SR49 | $6.46\times10^{-3}$ | 1.24 | 0.25 | 12.57 |
| SR50 | $9.18\times10^{-3}$ | 1.40 | 8.24 | 8.43 |
| SR51 | $1.25\times10^{-3}$ | 4.05 | $4.93\times10^{2}$ | 7.56 |
| SR52 | $5.06\times10^{-3}$ | 1.44 | $6.52\times10^{-2}$ | 13.42 |
| SR53 | $8.00\times10^{-2}$ | 1.30 | $2.15\times10^{-2}$ | 16.10 |
| SR54 | - | - | - | - |
| SR55 | $1.60\times10^{-4}$ | 2.00 | 8.72 | 8.22 |
| SR56 | $5.16\times10^{-6}$ | 8.15 | $2.12\times10^{2}$ | 11.17 |

**Supplementary Table** 2. **Kinetic parameters of nesprin SR40-SR56.** The zero-force unfolding rate ($k_u^0$), zero-force refolding rate ($k_f^0$), the unfolding transition distance ($x_u$) and the folding transtion rigid body length ($b^\star$) are listed. The $k_u^0$ and $x_u$ are obtained based on Bell's model fitting to the unfolding force distribution of the domains. The $k_f^0$ and $l^\star$ are obtained by fitting to the refolding force distribution of the domains based on Arrhenius Law. The cooperative unfolding/refolding domains are assigned with same parameters values. The parameters of those domains without clear unfolding/refolding dynamics are not obtained (indicated by '$-$').

#### Supplementary Note 1. Protein constructs sequence information

The detailed sequence information of the corresponding protein constructs are listed below:

**1.** AviTag- (linker)-(Nesprin-2G SR1-SR2)$^{299\text{-}474}$- (linker)-SpyTag001:

MHHHHHHGKPIPNPLLGLDSTENLYFQGIDPFT GLNDIFEAQKIEWHE GGGSG
KLAKVRDALVWLTLQEKRFQKMLKDSASETYCNKYHSLLSFMESLNEEKESF
IDVLSLKGRMGELNEDESRLRQGWTSLMHQVAAWRAQLDDALPSPLKETEA
WLKDIEGVVQEGVPTSQSYSEARTLIQGKLSSFKSLMGSFDYHSDVLMAFQSN
AEKSLPAVPPVKLEEMTRRINNEFGGGSGAHIVMVDAYKPTK

**2.** AviTag- (linker)-(Nesprin-2G SR3-SR4)$^{475\text{-}684}$- (linker)-SpyTag001:

MHHHHHHGKPIPNPLLGLDSTENLYFQGIDPFT GLNDIFEAQKIEWHE GGGSG
KLVLGKNFIPLLEFHDSKCSVLALLDEAKAKLDVWNGTYESKESVEVLLEDW
HKFTGEKKFLIQLDASFQKCEEMYKNSARECESIREEYMMLEKNVHSCRQYIH
NTKATLQRALMSWATFEEDLALLKASFDLTKKEQIKEVPVETLLQWNTKHTS
LNEVGSFLIGVSSREVAASISKELRRLNKRWRKFITKTPLLKLPLVKIQDQPPG
NEFGGGSGAHIVMVDAYKPTK

**3.** AviTag- (linker)-(Nesprin-2G SR5-SR7)$^{712\text{-}1050}$- (linker)-SpyTag001:

MHHHHHHGKPIPNPLLGLDSTENLYFQGIDPFT GLNDIFEAQKIEWHE GGGSG
KLEKQEVEDEESAGQLKVNEEVEGLIKQVTIWESQTKSILDLLQHGDHADGSS
ADTLQHLIAKGSVYEELLARTEDTLQMDVQSPSNLEPFQNVLRAGLQAKIQEA
KQGVQITMVELSAVLKNLSDEPLELDLGLKVEEAQKELEVSILRAEQLLGQRE
RPGGFLLKYKEALEILNTNSLAKYLRAVEELKRTVPGGAKLQLEEQSRVASAK
WEPLRHEISLYLQQLKIAIEEEKLRDNIARLEKQINKEKKLIRRGRTRGLRKEH
EACLSPESIKCQLEHHVGVLRVLCEELTSPEDQQELKRALRDYEQKIARLLKC
ASEIHTTLQSSQGGALEERSALEFGGGSGAHIVMVDAYKPTK

**4.** AviTag- (linker)-(Nesprin-2G SR8)$^{1062\text{-}1121}$- (linker)-SpyTag001:

MHHHHHHGKPIPNPLLGLDSTENLYFQGIDPFT GLNDIFEAQKIEWHE GFEID
KVWYDLDAKLGDIEFIKVNKGGGSGSGDATIPPPPPLPEGVGIPSPSSLPGGTAI
PPPPPGGGSGLEGEVPLEIPDNQLSTEKAMEPIKNFSQTSELKPQQEESIMEKE
GKDCSASLSDLQERYDTQRGLLEQHLQDSKSRVTSDFASEQERSSACLQSKLA
ELQVLLADTDAHWEKFEITSLNLRRLMSDAEKPVLNQERDLLKGNEQVLHGL
LNTGSGGSGSGEAGMPPPPPPLPGGPGIPPPPPFPGGPGIPPPPPGGGSGAHI
VMVDAYKPTK

**5.** AviTag- (linker)-(Nesprin-2G SR9-SR10)$^{1262\text{-}1409}$- (linker)-SpyTag001:

MHHHHHHGKPIPNPLLGLDSTENLYFQGIDPFT GLNDIFEAQKIEWHE GGGSG
KLDIRDSIAKQIRVCTSLEEPSNSVPRELHTLDQCAIQDIVLKCRLQLETMNQK
VEMREDALDALEGFLASLRAAKLSAELPADRPAPKAPEVLSEDILLMKEKAG
PLDERLRTLGINIKDAEGGENTTCERLVGALSVNLVAMDGQSKEEGPPEDKK
LLEACSSKNLELFKNIQDLQNQISKIGLKDPTAPAVKHRKKSLLRLDKDLDGLE
EEKVRIQKIAGSLPRFKDGSEKNVIQQCEDTAALWESTKASVTESLEQCGSAL
ELLRQYQNIKNNLTALIQKEEGIISQQASYMGKDNLKKKIAEIETVKEEFSDHL

EVVDKINQICKNLQYHLNKMKTFEDPPFEKEANAIVDRWLDINEKTEEYGEN
LGRALALWDKLFIIKNNIDEWTEQILGKAESHELTEEDRGRLKEELKVLEEQS
AEFSRRVADIQSLLQSNEKPLELQVMESSVLSKMKDVKTHVAGGSNSYAPSGS
TAELREDLDQAKTQMGMTESLLNALSPSDSLEIFTKLEEIQQQIFQQKHSMTV
LENQIGCLTPELSELKRQYASVSNLFNTKKNALQDHFATFLNEQCKNFNDWF
SNVKTNLQECFEPPETKLSLEQRLQKLSDFLTLGGGNSKIQQVETVLQHVKML
LPKAHVKELDSWLRSQELELENMESICQARAGELNNSFQQLLRLEDDCRSLSK
WLTNQEENWGKMEVSGERMDLFSQALTRKREQFETVAQLSDSLKEHGLTEG
EETIKESTHLIDRYQALWRQLHEIEEEDKLPAAEDQSFNDLADDVIHWIKEIKE
SLMALNSSEGKMPLEERIQKIKEIIALKPEGDAKIQMVMRQAEHCEAPLAQET
FTDLSNQWDSTLHLANTYLSHQEKLVLEGEKYLQSKEDLRLMLTELKKQQEA
GFALQPGLPEKQAQLKIYKKFLQKAQDLTSLLEELKSQGNYLLECTKNPSFSE
EPWLEVKHLHESLLQQLQDSVQKLEGHVQEHSSYQVCLTDLSSTLDDISKEYF
SLCDGSKDQIMAKERMQKLQELESRLRFQGGALKKASALAKSIKQNTSSVGQ
KIIKDDIRSLKYKQKDLENRIESAKQETENGLNSIEFGGGSGAHIVMVDAYKPT
K

**6**. AviTag- (linker)-(Nesprin-2G SR11-SR13)$^{1410\text{-}1728}$- (linker)-SpyTag001:

MHHHHHHGKPIPNPLLGLDSTENLYFQGIDPFTGLNDIFEAQKIEWHEGGGSG
KLEGPPEDKKLLEACSSKNLELFKNIQDLQNQISKIGLKDPTAPAVKHRKKSLL
RLDKDLDGLEEEKVRIQKIAGSLPRFKDGSEKNVIQQCEDTAALWESTKASVT
ESLEQCGSALELLRQYQNIKNNLTALIQKEEGIISQQASYMGKDNLKKKIAEIE
TVKEEFSDHLEVVDKINQICKNLQYHLNKMKTFEDPPFEKEANAIVDRWLDIN
EKTEEYGENLGRALALWDKLFIIKNNIDEWTEQILGKAESHELTEEDRGRLKE
ELKVLEEQSAEFSRRVADIQSLLQSNEKPLELQVMESSVLSKMKDVKTHVAGG
SEFGGGSGAHIVMVDAYKPTK

**7**. AviTag- (linker)-(Nesprin-2G SR14-SR16)$^{1729\text{-}1820}$- (linker)-SpyTag001:

MHHHHHHGKPIPNPLLGLDSTENLYFQGIDPFTGLNDIFEAQKIEWHEGGGSG
KLNSYAPSGSTAELREDLDQAKTQMGMTESLLNALSPSDSLEIFTKLEEIQQQI
FQQKHSMTVLENQIGCLTPELSELKRQYASVSNLFNTKKNALQDHFATFLNE
QCKNFNDWFSNVKTNLQECFEPPETKLSLEQRLQKLSDFLTLGGGNSKIQQV
ETVLQHVKMLLPKAHVKELDSWLRSQELELENMESICQARAGELNNSFQQLL
RLEDDCRSLSKWLTNQEENWGKMEVSGERMDLFSQALTRKREQFETVAQLS
DSLKEHGLTEGEETIKESTHLIDRYQALWRQLHEIEEEDEFGGGSGAHIVMVD
AYKPTK

**8**. AviTag- (linker)-(Nesprin-2G SR17-SR19)$^{2027\text{-}2350}$- (linker)-SpyTag001:

MHHHHHHGKPIPNPLLGLDSTENLYFQGIDPFTGLNDIFEAQKIEWHEGGGSG
KLKLPAAEDQSFNDLADDVIHWIKEIKESLMALNSSEGKMPLEERIQKIKEIIAL
KPEGDAKIQMVMRQAEHCEAPLAQETFTDLSNQWDSTLHLANTYLSHQEKL
VLEGEKYLQSKEDLRLMLTELKKQQEAGFALQPGLPEKQAQLKIYKKFLQKA
QDLTSLLEELKSQGNYLLECTKNPSFSEEPWLEVKHLHESLLQQLQDSVQKLE
GHVQEHSSYQVCLTDLSSTLDDISKEYFSLCDGSKDQIMAKERMQKLQELESR
LRFQGGALKKASALAKSIKQNTSSVGQKIIKDDIRSLKYKQKDLENRIESAKQE
TENGLNSIEFGGGSGAHIVMVDAYKPTK

**9**. AviTag- (linker)-(Nesprin-2G SR20-SR21)$^{2422\text{-}2610}$- (linker)-SpyTag001:

MHHHHHHGKPIPNPLLGLDSTENLYFQGIDPFT GLNDIFEAQKIEWHE GGGSG
KLDEREVNELQNQPLELDIMLRNEQLKGMEELSTHLEARRAAIELLEQSQHLN
QTEEQALVLPAARPSVCHLGSLQQELHTLKKTKERQYGLLSGFQDQLVMAEA
SLNTSLAEVESLKIGSLDSATYLGKIKKFLGSVENQQGSLSKLRTEWAHLSSLL
AAADQKLVESQMKHLEHGWELVEQLAHRKCFQEFGGGSGAHIVMVDAYKP
TK

**10**. AviTag- (linker)-(Nesprin-2G SR22-SR23)$^{2611\text{-}2821}$- (linker)-SpyTag001:

MHHHHHHGKPIPNPLLGLDSTENLYFQGIDPFT GLNDIFEAQKIEWHE GGGSG
KLQATEHSELTCLLEKLQDLKVSLHQQQQRLTLSLNSPGQQAAIVDMVTPAA
ELQAIKCEFSGLKWQAELHMKRLWGEKDKKTLEDAINNLNKQMEALEPLNR
EVENRIKKCELQNRIKETLSWVKNTMAELVVPIALLPDNILSQIRKCKLIHDGIL
GNQQAVELLVEEVRGITPSLAPCEGDGLNALLEDLQSQHQALLLKSTERSQQL
ELEFGGGSGAHIVMVDAYKPTK

**11**. AviTag- (linker)-(Nesprin-2G SR24-SR25)$^{2822\text{-}3027}$- (linker)-SpyTag001:

MHHHHHHGKPIPNPLLGLDSTENLYFQGIDPFT GLNDIFEAQKIEWHE GGGSG
KLKLEEKSKLFAIIGKVQLTLEESETLMSPTGDRASTEAELERRLAILKASQQQ
LQDTESALSAHLQELTNAYKDANVFERLFLDDQLKNLKARTNRTQRFLQNNG
SELKQKMESYREFHDKAAVLQKEAECILHGGLLPLRQELEQDAKEQLGNLRD
KLAAIRGSLSQVLTSEEVFDTIGLSWDGSLLARLQTQVLEREREVEGKIKEFGG
GSGAHIVMVDAYKPTK

**12**. AviTag- (linker)-(Nesprin-2G SR26-SR27)$^{3028\text{-}3239}$- (linker)-SpyTag001:

MHHHHHHGKPIPNPLLGLDSTENLYFQGIDPFT GLNDIFEAQKIEWHE GGGSG
KLQLDTFLIARDRHQASISKIRAVDLQIKKGAESLLKVPSMSPESTLLNAQTLIQ
KIEKSKRLRDEIIRKLSKNEAFDDSFKESEMQRLKLCAEENSRLQEALQNMLLE
LQPREMGEKEFREKLENALHVLKQIQSRLQQPLCVNLGVQHIQHEKETWEAF
GEQVEAEMCGLRAVRITEEQREENDSGTGGMEAKLRDIEGLHMELSKSISLRA
EFGGGSGAHIVMVDAYKPTK

**13**. AviTag- (linker)-(Nesprin-2G SR28-SR29)$^{3240\text{-}3456}$- (linker)-SpyTag001:

MHHHHHHGKPIPNPLLGLDSTENLYFQGIDPFT GLNDIFEAQKIEWHE GGGSG
KLDVLNDAYDSANRYDELVAGALRIITSLEATLLSYRVDLHNPQKTLELAHLK
QEELQSSVADLRSLTETLGAISSPEAKEQLRCTLEVLAAKNSALKAGLEAQEAE
EERCLENYKCFRKMKEEICSRLRKMEMDLGQSIFPLPRSYKEALARLEQSKAL
TSNLLSTKEDLVKLRQLLRHLRCRSTENDATCALGVASALWEKWLSLLEAAR
EWQQWGGEFGGGSGAHIVMVDAYKPTK

**14**. AviTag- (linker)-(Nesprin-2G SR30-SR31)$^{3457\text{-}3669}$- (linker)-SpyTag001:

MHHHHHHGKPIPNPLLGLDSTENLYFQGIDPFT GLNDIFEAQKIEWHE GGGSG
KLELKREWKFISEEIEREAIILETLQEDLPEISKTNAAPTEELWQLLDSLCQHQE
SVEKQQLLLALLLQRVRSIQNIPEGTETGETIPALQEIGSMQERCDRLLHTTRK
NKDLVQAEIQAQQSFLKEIKDVKRVFEQISTSFPNLAPEGHPERAEQFEELRSI
LQKGKLSFENIMEKLRIKYSEMYSIVPAEIGSQVEECRSALEDAEEKMSSEVEF
GGGSGAHIVMVDAYKPTK

**15**. AviTag- (linker)-(Nesprin-2G SR32-SR33)$^{3670\text{-}3870}$- (linker)-SpyTag001:

MHHHHHHGKPIPNPLLGLDSTENLYFQGIDPFT GLNDIFEAQKIEWHE GGGSG
KLSKSSPSSIMRRKIERINNGLHCVEKMLQQKSRNIEEAHEIQKKIWDELDLWH
SKLNELDSEVQDFVEQDPGQAQEWMDNLMAPFQQHQQVSQRAESRTSQLNK
ATIKMEEYNDLLKSTEVWIEKTSCLLANPACYDSSRTLSHRASTLQMALEDSE
QKHSLLHSIFTDLEDLSIIFETDDLIQTIHELSDQVAALQQQIMEEFGGGSGAHIV
MVDAYKPTK

**16**. AviTag- (linker)-(Nesprin-2G SR34-SR35)$^{3871\text{-}4074}$- (linker)-SpyTag001:

MHHHHHHGKPIPNPLLGLDSTENLYFQGIDPFT GLNDIFEAQKIEWHE GGGSG
KLALPHVQQVADDVVAIESEVKAMEKKVAKIKAILLSKEIFDFPPEEHLKHGE
VILENIHPMKKTIAEIMTYQVELRLPQTGTKPLPVFQRTSQLLQDVKLLENVT
QEQNELLKVVIKQTAECDEEIDSLKQMLTNYSAEISPEHVSQNQVADLPSLQG
EMERLEKQILNLNQKKEDLLVDLKTAVLNLHEHLKQEQQEVGDKPSAEFGGG
SGAHIVMVDAYKPTK

**17**. AviTag- (linker)-(Nesprin-2G SR36)$^{4218\text{-}4337}$- (linker)-SpyTag001:

MHHHHHHGKPIPNPLLGLDSTENLYFQGIDPFT GLNDIFEAQKIEWHE GFEID
KVWYDLDAKLGDIEFIKVNKGGGSGSGDATIPPPPPLPEGVGIPSPSSLPGGTAI
PPPPPGGGSGLERSRPRPADILRVCKTQVAKLELWLQQANVAFEPETVDADM
QQVVEEELAGCQAMLTEIEYKVASLLETCKDQGLGDCGTTQQEAEALSWKL
KTVKCNLEKVQMVLQEKFSEDQHPSTLKKGSGGSGSGEAGMPPPPPPLPGG
PGIPPPPPPFPGGPGIPPPPPGGGSGAHIVMVDAYKPTK

**18**. AviTag- (linker)-(Nesprin-2G SR37-SR38)$^{4507\text{-}4714}$- (linker)-SpyTag001:

MHHHHHHGKPIPNPLLGLDSTENLYFQGIDPFT GLNDIFEAQKIEWHE GGGSG
KLSMTEETYISLDKRLFELFLSLSRCLGSVEGLLQRPGLLREDACAQQVFFQKL
ALELKKLYLALGDKKDDFLKAVTWPGKEATLLPECIDALTVSLESVQSRAAW
RDASLKAGLEHSRSYQNEVKRLYSQLIKKKTALQQSLNEISGQSISKQLQKADV
HTAELQNSEKQVAKLRDEGERLRFPHGLLQDVYKLEDVLDSMWGILRARYEF
GGGSGAHIVMVDAYKPTK

**19**. AviTag- (linker)-(Nesprin-2G SR39-SR40)$^{4715\text{-}4929}$- (linker)-SpyTag001:

MHHHHHHGKPIPNPLLGLDSTENLYFQGIDPFT GLNDIFEAQKIEWHE GGGSG
KLLELSSPFLSKSLQTLLQGMAELVSIGKGKLAADPLQHAKSKAALQAQLQDH
KAFFQKLVADMLLIQTYSATMFPPSLQKGEGFGAEQVAEVRALEEEACLRGA
QLQSMLQKWEEFDDNYASLEKDLEALISSLPSVSLVEETEERLLERISFYQQIK
RNIDGKHARLCQTLNEGRQLAASVSCPEPEGQIARLEEQWLSLNKRIDQELHR
LQTLLEFGGGSGAHIVMVDAYKPTK

**20**. AviTag- (linker)-(Nesprin-2G SR41-SR42)$^{4930\text{-}5150}$- (linker)-SpyTag001:

MHHHHHHGKPIPNPLLGLDSTENLYFQGIDPFT GLNDIFEAQKIEWHE GGGSG
KLKHLLSYSRDSDELTRWLETSQQTLNYWKEQSLNVSQDLNTIRSNIDRFFKF
SKEVDERSSLKSAVMSTGNQLLHLKEADTATLRASLAQFEQKWTVLITQLPDI
QEKLHQLQMEKLPSREAISEMISWMNAVEPQAAGKDTELSKSSASQVKHLLQ
KLKEFRMEMDYKQWVVDFVNQSLLQLSTCDVESKRYERTEFAEHLGEMNRQ
WQRVHGTLNRKIQHEFGGGSGAHIVMVDAYKPTK

**21.** AviTag- (linker)-(Nesprin-2G SR43-SR44)[5151-5377]- (linker)-SpyTag001:

MHHHHHHGKPIPNPLLGLDSTENLYFQGIDPFT GLNDIFEAQKIEWHE GGGSG
KLLEQLLESITENENKVQNLNSWLEAQEERLKMLQKPESAVSMEKLLLDCQDI
ENQLALKSKALDELRQSSLTMDGGDVPLLEDMASGIVELFQKKNNVTSQVHQ
LRASVQSVLQEWKACDKLYDEATMRTTQLTYSMEHSKPAVLSLQALACQVQ
NLEALQDEAENGERSWEKLQEVIGRLKASCPSMAGIIEEKCQDAHSRWTQVN
QDLADQLQEARGQLQLWKAPHEFGGGSGAHIVMVDAYKPTK

**22.** AviTag- (linker)-(Nesprin-2G SR45-SR46)[5378-5576]- (linker)-SpyTag001:

MHHHHHHGKPIPNPLLGLDSTENLYFQGIDPFT GLNDIFEAQKIEWHE GGGSG
KLNAHAEAAAWLQQQEAKFQQLANTNLSGDNLADILPRALKDIKGLQSDLQK
TKEAFLENSTLSDQLPQPEERSTPGLHSGQRHSLQTAAYLEKMLLAKSNEFEI
VLAQFKDFTDRLAYSKDLIVHKEENLNKLYHEEKEEVPDLFLNHVLALTAQSP
DIERLNEESLRLPLSDVTIKTLQSLNRQWIRATATALDHYSELEFGGGSGAHIV
MVDAYKPTK

**23.** AviTag- (linker)-(Nesprin-2G SR47-SR48)[5577-5786]- (linker)-SpyTag001:

MHHHHHHGKPIPNPLLGLDSTENLYFQGIDPFT GLNDIFEAQKIEWHE GGGSG
KLQGNGLNEKFLHYCERWIQVLEKIQESLSVEVAHSLPALLEQQKTYEILEAE
VSTNQAVADAYVTQSLQLLDTAEIEKRPEFVSEFSKLSDQWQRAARGVRQRK
CDISRLVTQWRFFTTSVEDLLRFLADTSQLLSAVKEQDCYSLCQTRRLVHELK
SKEIHLQRWRTTYALALEAGEKLRNTPSPETREFVDGQISRLQESWKDTELSL
GEFGGGSGAHIVMVDAYKPTK

**24.** AviTag- (linker)-(Nesprin-2G SR49-SR51)[5787-6122]- (linker)-SpyTag001:

MHHHHHHGKPIPNPLLGLDSTENLYFQGIDPFT GLNDIFEAQKIEWHE GFEID
KVWYDLDAKLGDIEFIKVNKGGGSGSGDATIPPPPPLPEGVGIPSPSSLPGGTAI
PPPPPGGGSGLEEVISRLQSTAETWDQCKKKIKKLKKRLQALKAQSEDPLPEL
HEALHEEKELIKEVEKSLANWTHSLKELQTMKADLSQHILAEDVTVLKEQIQL
LHRQWEDLCLRVAIRKQEIEDRLNSWIVFNEKNKELCAWLVQMENKVLQTA
DVSIEEMIEKLQKDCMEEISLFTENKLQLKQMGDQLIKASSKAKAAELEEKLSK
INDRWQHLFDVIGSRVKKLKETFAFIQQLDKNMSNLRTWLARIESELSKPVVY
DVCDNQEIQKRLAEQQDLQRDIEQHSAGVESVFNICDVLLHDSDACANETECD
SIQQTTRSLDRRWRNICAMSMERRMKIEETWGSGGSGSGEAGMPPPPPPLPG
GPGIPPPPPFPGGPGIPPPPPGGGSGAHIVMVDAYKPTK

**25.** AviTag- (linker)-(Nesprin-2G SR52-SR53)[6110-6362]- (linker)-SpyTag001:

MHHHHHHGKPIPNPLLGLDSTENLYFQGIDPFT GLNDIFEAQKIEWHE GGGSG
KLMSMERRMKIEETWRLWQKFLDDYSRFEDWLKSAERTAACPNSSEVLYTN
AKEELKRFEAFQRQIHERLTQLELINKQYRRLARENRTDTASKLKQMVHEGN
QRWDNLQKRVTAILRRLRYFTNQREEFEGTRESILVWLTEMDLQLTNVEHFS
ESDAEDKMRQLNGFQQEITLNTNKIDQLIVFGEQLIQKSEPLDAVLIEDELEEL
HRYCQEVFGRVSRFHRRLTSHTPGLDDEKEASENETDIEDPREIQAEFGGGSG
AHIVMVDAYKPTK

**26.** AviTag- (linker)-(Nesprin-2G SR54-SR56)[6433-6822]- (linker)-SpyTag001:

MHHHHHHGKPIPNPLLGLDSTENLYFQGIDPFT GLNDIFEAQKIEWHE GGGSG KLSGKSISEGHPWHVPDSPSHSKHHYKHMEGDRTEAPVPTDASTPFKSDYVK LLLRQGTDDSKEGLKEAQQEDEQLATLTGQQPGAFDRWELIQAQELHSKLRL KQTVQQLKSDIGSIAAWLGKTEAELEALKLAEPPSDIQEIALRVKRLQEILKAF DTYKALMVSVNVSHKEYLPSQSPEATELQNRLHQLSLSWDSVQGVLDSWRGD LRQSLMQCQDFHQLSQDLLLWLATAESRRQKAHVTSPEADRQVLLECQKDL MRLEKELVARQPQVSSLREISSSLLVKGQGEDYIEAEEKVHVIEKKLKQLQEQ VAQDLMSLQRSLDPDASLTSFDEVDSGEQLPAAFAKFGVEEEEEEEETDSRMP HLDSPGSSQPRRSFLSRVIEFGGGSGAHIVMVDAYKPTK

**27**. AviTag- (linker)-(Nesprin-2G SR54-SR56)$^{6110\text{-}6822}$- (linker)-SpyTag001:

MHHHHHHGKPIPNPLLGLDSTENLYFQGIDPFT GLNDIFEAQKIEWHE GGGSG KLMSMERRMKIEETWRLWQKFLDDYSRFEDWLKSAERTAACPNSSEVLYTN AKEELKRFEAFQRQIHERLTQLELINKQYRRLARENRTDTASKLKQMVHEGN QRWDNLQKRVTAILRRLRYFTNQREEFEGTRESILVWLTEMDLQLTNVEHFS ESDAEDKMRQLNGFQQEITLNTNKIDQLIVFGEQLIQKSEPLDAVLIEDELEEL HRYCQEVFGRVSRFHRRLTSHTPGLDDEKEASENETDIEDPREIQADSWRKR RESEEPTSPQSLCHLVPPALGHERSGCETPVSVDSIPLEWDHTGDVGGSSSHE DDEEGPFYSALSDVEIPENPEAYLKMTTKSLQASSGKSISEGHPWHVPDSPSHS KHHYKHMEGDRTEAPVPTDASTPFKSDYVKLLLRQGTDDSKEGLKEAQQED EQLATLTGQQPGAFDRWELIQAQELHSKLRLKQTVQQLKSDIGSIAAWLGKT EAELEALKLAEPPSDIQEIALRVKRLQEILKAFDTYKALMVSVNVSHKEYLPSQ SPEATELQNRLHQLSLSWDSVQGVLDSWRGDLRQSLMQCQDFHQLSQDLLLW LATAESRRQKAHVTSPEADRQVLLECQKDLMRLEKELVARQPQVSSLREISSS LLVKGQGEDYIEAEEKVHVIEKKLKQLQEQVAQDLMSLQRSLDPDASLTSFDE VDSGEQLPAAFAKFGVEEEEEEEETDSRMPHLDSPGSSQPRRSFLSRVIEFGG GSGAHIVMVDAYKPTK

**28**. AviTag- (linker)-(Nesprin-2G SR09-SR19)$^{1262\text{-}2350}$- (linker)-SpyTag001:

MHHHHHHGKPIPNPLLGLDSTENLYFQGIDPFT GLNDIFEAQKIEWHE GGGSG KLMSMERRMKIEETWRLWQKFLDDYSRFEDWLKSAERTAACPNSSEVLYTN AKEELKRFEAFQRQIHERLTQLELINKQYRRLARENRTDTASKLKQMVHEGN QRWDNLQKRVTAILRRLRYFTNQREEFEGTRESILVWLTEMDLQLTNVEHFS ESDAEDKMRQLNGFQQEITLNTNKIDQLIVFGEQLIQKSEPLDAVLIEDELEEL HRYCQEVFGRVSRFHRRLTSHTPGLDDEKEASENETDIEDPREIQADSWRKR RESEEPTSPQSLCHLVPPALGHERSGCETPVSVDSIPLEWDHTGDVGGSSSHE DDEEGPFYSALSDVEIPENPEAYLKMTTKSLQASSGKSISEGHPWHVPDSPSHS KHHYKHMEGDRTEAPVPTDASTPFKSDYVKLLLRQGTDDSKEGLKEAQQED EQLATLTGQQPGAFDRWELIQAQELHSKLRLKQTVQQLKSDIGSIAAWLGKT EAELEALKLAEPPSDIQEIALRVKRLQEILKAFDTYKALMVSVNVSHKEYLPSQ SPEATELQNRLHQLSLSWDSVQGVLDSWRGDLRQSLMQCQDFHQLSQDLLLW LATAESRRQKAHVTSPEADRQVLLECQKDLMRLEKELVARQPQVSSLREISSS LLVKGQGEDYIEAEEKVHVIEKKLKQLQEQVAQDLMSLQRSLDPDASLTSFDE VDSGEQLPAAFAKFGVEEEEEEEETDSRMPHLDSPGSSQPRRSFLSRVIEFGG GSGAHIVMVDAYKPTK

**Supplementary Note 2. Protein expression and purification protocol**

Plasmids containing the target protein domains were co-transformed with a BirA plasmid into Escherichia coli BL21 (DE3) via heat shock (42 °C for 30 seconds), followed by overnight incubation on an LB agar plate at 37 °C for at least 12 hours. A single colony was selected and cultured in 5 mL of LB broth at 37 °C with shaking at 250 rpm overnight (12–14 hours). Subsequently, 4 mL of the overnight culture was transferred into 400 mL of fresh LB broth (1:100 dilution) and incubated at 37 °C with shaking at 250 rpm for 2–3 hours, until the optical density (OD) reached 0.4–0.6. D-Biotin and IPTG were then added to final concentrations of 50 $\mu$M and 0.4 mM, respectively. The culture was allowed to grow at 20 °C with shaking at 250 rpm overnight (approximately 16 hours). Afterward, the cells were harvested by centrifugation at 4000 rpm at room temperature, and the pellet was resuspended in bacterial lysis buffer consisting of 50 mM Tris (pH 7.5), 500 mM NaCl, 10% glycerol, and 20 mM imidazole. Cell lysis was carried out by sonication, and the resulting lysate was incubated with Ni-NTA resin for approximately 3 hours at 4 °C with gentle shaking. The protein was then eluted using a buffer containing 50 mM Tris (pH 7.5), 500 mM NaCl, 10% glycerol, 1 mM $\beta$-Mercaptoethanol, and 200 mM imidazole. Imidazole was removed through buffer exchange by dialysis or using the ÄKTA system. The final protein concentration was approximately 1 mg mL$^{-1}$ in storage buffer (50 mM Tris, pH 7.5, 500 mM NaCl, 5 mM DTT, and 10% glycerol). Protein aliquots were snap-frozen in liquid nitrogen and stored at -80 °C.
